# The Calcineurin-TFEB-p62 Pathway Mediates the Activation of Cardiac Macroautophagy by Proteasomal Malfunction

**DOI:** 10.1101/2019.12.27.889519

**Authors:** Bo Pan, Nirmal Parajuli, Zongwen Tian, Jie Li, Penglong Wu, Megan T. Lewno, Lynn Bedford, R. John Mayer, Jing Fang, Jinbao Liu, Taixing Cui, Huabo Su, Xuejun Wang

**Author notes:** These authors contributed equally (Bo Pan, Nirmal Parajuli, Zongwen Tian). **Address correspondence to:** Dr. Xuejun Wang, Division of Basic Biomedical Science, Sanford School of Medicine of the University of South Dakota, Vermillion, SD 57069, USA; or Dr. Huabo Su, Vascular Biology Center and Department of Pharmacology and Toxicology, Medical College of Georgia, Augusta University; Augusta, GA, USA.

## Abstract

**Rationale:** The ubiquitin-proteasome system (UPS) and the autophagic-lysosomal pathway (ALP) are pivotal to proteostasis. Targeting these pathways is emerging as an attractive strategy for treating cancer. However, a significant proportion of patients who receive a proteasome inhibitor-containing regime for example, show cardiotoxicity. Moreover, UPS and ALP defects are implicated in the pathogenesis of a large subset of heart disease. Hence, a better understanding of the cross-talk between the two catabolic pathways should help advance cardiac pathophysiology and medicine.

**Objective:** Systemic pharmacological proteasome inhibition (PSMI) was shown to increase p62/SQSTM1 expression and induce myocardial macroautophagy. The present study investigates whether cardiomyocyte-restricted PSMI activates myocardial ALP and, more importantly, how proteasome malfunction activates the ALP in the heart.

**Methods and Results:** Myocardial macroautophagy, transcription factor EB (TFEB) expression and activity, and p62 expression were markedly increased in mice with either cardiomyocyte-restricted ablation of *Psmc1* (a 19S proteasome subunit gene) or pharmacological PSMI. In cultured cardiomyocytes, PSMI-induced increases in TFEB activation and p62 expression were blunted markedly by calcineurin inhibition (cyclosporine A) and by siRNA-mediated *Molcn1* silence. PSMI induced remarkable increases in myocardial autophagic flux in wild type mice but not p62 null mice. In cultured wild type, but not p62-null, mouse cardiomyocytes, PSMI induced increases in LC3-II flux and in the lysosomal removal of ubiquitinated proteins. Myocardial TFEB activation by PSMI as reflected by TFEB nuclear localization and target gene expression was strikingly less in p62 null mice compared with wild type mice.

**Conclusions:** (1) The activation of cardiac macroautophagy by proteasomal malfunction is mediated by the Mocln1-calcineurin-TFEB-p62 pathway; (2) both Mocln1 and p62 form a feed-forward loop with TFEB during TFEB activation by proteasome malfunction; and (3) targeting the Mcoln1-calcineurin-TFEB-p62 pathway may provide new means to intervene cardiac ALP activation in a proteasome malfunction setting.

## INTRODUCTION

The ubiquitin-proteasome system (UPS) is responsible for the degradation of most proteins in the cell. The UPS is pivotal to both protein quality control and the regulatory degradation of normal proteins essential to virtually all cellular processes.^1^ The autophagic-lysosomal pathway (ALP) also plays a crucial role in intracellular quality control via removal of aberrant protein aggregates and defective or surplus organelles, in addition to provision of fuels by self-digesting a portion of cytoplasm during energy crisis.^2^ The UPS and ALP were historically believed to function in parallel in the cell but emerging evidence suggests the two pathways do cross-talk although the molecular mechanisms underlying the crosstalk remain poorly understood.^3^

In UPS-mediated protein degradation, the 26S proteasome carries out the final proteolytic step of poly-ubiquitinated proteins and, as indicated by emerging evidence, its functioning is highly regulated and often constitutes the rate–limiting step.^4^ Studies on human myocardium with cardiomyopathies and end-stage heart failure have yielded compelling evidence that cardiac proteasome impairment occurs in a large subset of heart disease during progression to heart failure and may play a major pathogenic role therein.^5, 6^ Studies using animal models have established the necessity of proteasome impairment or functional insufficiency in both rare and common forms of cardiac disorders such as cardiac proteinopathy,^7^ myocardial ischemia/reperfusion injury,^7-9^ pressure-overloaded cardiac hypertrophy and failure,^10, 11^ as well as diabetic cardiomyopathy.^12^ Therefore, beyond search for measures to improve cardiac proteasome functioning, a better understanding of how cardiomyocytes and the heart respond to proteasome impairment will help identify potential strategies to assist the heart in maintaining proteostasis. Prior studies have shown that proteasome inhibition (PSMI) with pharmacological agents increases myocardial macroautophagy (hereafter referred to as autophagy);^13^ and both autophagy and p62/SQSTM1 in cardiomyocytes are markedly increased in the compensatory stage of cardiac proteinopathy. The latter exemplifies heart disease with increased proteotoxic stress and is associated with proteasome impairment.^14-17^ However, the molecular mechanisms governing the activation of autophagy by proteasome malfunction and the role of p62 upregulation in the induction of cardiac autophagy by PSMI remain undefined.

Transcription factor EB (TFEB) has emerged as a master regulator of lysosomal genesis, the ALP, and catabolism. At baseline, TFEB is phosphorylated at multiple residues by kinases including the all-important mechanistic target of rapamycin complex 1 (mTORC1) and possibly extracellular signal-regulated kinase 2 (also known as MAPK1) and MAP4K3.^18^ Bound by cytosolic chaperone 14-3-3, the phosphorylated TFEB is segregated in the cytoplasm where the TFEB proteins or, at least, a large fraction of them are localized on the membrane of lysosomes. During starvation or lysosomal stress, mTORC1 is inactivated and stops phosphorylating TFEB while TFEB is dephosphorylated by calcineurin, allowing TFEB nuclear translocation. Here the activation of calcineurin is induced by the release of lysosomal Ca^2+^ via the Ca^2+^ channel mucolipin 1 (MCOLN1).^19^ In the nucleus, TFEB directly binds to a common 10-base E box-like palindromic sequence, referred to as the coordinated lysosomal expression and regulation (CLEAR) element, and thereby activates the transcription of an entire network of genes harboring the CLEAR motif. This network is known as the CLEAR network which consists of genes involved in lysosomal genesis, autophagosome formation and even mitochondrial biogenesis.^20^ The role of TFEB activation in cardiac pathophysiology has begun to unveil;^15, 21-24^ however, it remains untested how TFEB participates in the crosstalk between the UPS and ALP, especially in bridging proteasome malfunction and autophagy activation. Hence, we performed the present study to address these critical questions.

After providing the first in vivo demonstration that genetic PSMI activates autophagy in mammalian hearts and cardiomyocytes, the present study has identified that both TFEB activation and p62 upregulation were induced by both genetic and pharmacological PSMI in the heart and cardiomyocytes, established that TFEB activation by PSMI is calcineurin- and Mcoln1-dependent, and demonstrated that p62 is required for induction of autophagy by PSMI and for lysosomal removal of ubiquitinated proteins in cardiomyocytes with proteasome malfunction. Taken together, the present study identifies the Mcolon1-calcineurin-TFEB-p62 pathway for proteasome malfunction to induce autophagy in the heart, which yields new mechanistic insight into the crosstalk between the UPS and ALP and provides potentially new therapeutic targets for modulating cardiac proteostasis.

## MATERIALS AND METHODS

The authors declare that all supporting data are available within the article and its online supplementary files.

### Animal models

The mice harboring the *Psmc1* floxed allele (Psmc1^f/f^),^25^ the transgenic (tg) mouse expressing the *Cre* recombinase driven by the mouse alpha myosin heavy chain promoter [B6.FVB-Tg(Myh6-cre)2182Mds/J; also known as αMyHC-Cre],^26^ p62 null (p62^-/-^) mice,^27^ GFPdgn tg mice,^28^ and the GFP-LC3 tg mice,^29^ were previously described. All mice were converted to C57BL/6J background before being used for this current study. In Psmc1^f/f^ mice, exon 2 and 3 are flanked by lox*P* sites. Psmc1^f/f^ mice were cross-bred with αMyHC-Cre::Psmc1^f/+^ mice to generate homozygous Psmc1^CKO^ (αMyHC-Cre::Psmc1^f/f^) mice, with their littermates of other genotypes used as controls. All protocols for animal use and care were approved by the University of South Dakota Institutional Animal Care and Use of Committee and conform to the NIH Guide for the Care and Use of Laboratory Animals. Animals used for this study had ad lib access to food and water and were housed in specific pathogen free control rooms with optimum temperature (22-24°C) and a 12-hour light and dark cycle.

### Neonatal rat or mouse cardiomyocyte cultures and genetic manipulations

Ventricular cardiomyocytes isolated from rats or mice at postnatal day 2 or day 1 respectively using the Neonatal Cardiomyocyte isolation System (Worthington) were cultured as previously reported.^22^ To induce the deletion of *Psmc1*, cardiomyocytes isolated from *Psmc1f/f* mice were infected with 20 M.O.I of adenoviruses expressing *Cre* (Ad-Cre) or β-galactosidase (Ad-β-Gal, as viral infection control).

The small interference RNA (siRNA) specific for rat Mcoln1 (Cat. #: 4390815) was purchased from ThemoFisher Scientific. The siRNA targeting rat Psmc1 included Rn_RGD:621097_1 FlexiTube siRNA (Qiagen, Cat# SI02002427) and Rn_RGD:621097_2 FlexiTube siRNA (Qiagen, Cat# SI02002434). The siRNA targeting luciferase (5’-AACGTACGCGGAATACTTCGA-3’) was used as the control siRNA. Lipofectamine™ 2000 transfection reagent (Invitrogen) was used for siRNA transfection to the cultured neonatal rat ventricular myocytes (NRVMs) at 72 hours after the cells were plated.^14^

### Protein extraction and western blot analyses

The extraction of total proteins from ventricular myocardial samples was done using a buffer containing (50 mM Tris-HCl at pH 6.8 containing 2% SDS, 10% glycerol and a complete protease inhibitor cocktail). A cocktail of protease inhibitors (#P-1540, AG. Scientific, San Diego, CA) were added to the extraction buffer to inhibit protein degradation. A similar protocol was used for the extraction of total proteins form cultured cardiomyocytes as well.^22^ Protein concentration was determined using bicinchoninic acid reagents (#23225, ThemoFisher Scientific, Waltham, MA). Equal amounts of proteins were fractionated via 8 ∼ 14% sodium dodecyl sulfate polyacrylamide gel electrophoresis (SDS-PAGE) and the separated proteins were transferred onto polyvinylidene difluoride (PVDF) membrane using a Trans-blot apparatus (Bio-Rad, Hercules, CA). The PVDF membranes were then sequentially subject to blocking, incubation with the primary antibodies against the protein of interest, washing with TBS-T buffer to remove unbound primary antibodies, incubation with horseradish peroxidase (HRP) conjugated secondary antibodies (Santa Cruz Biotechnology), and washing to remove unbound antibodies. The secondary antibodies bound to the PVDF membrane were then detected using enhanced chemiluminescence reagents (GE Healthcare, NJ); the chemiluminescence was digitally imaged and analyzed with the ChemiDocTM MP imaging system and associated software (Bio-Rad, Hercules, CA 94547) as we previously reported.^15^ The antibodies used are described in **Supplementary Table 1**. For loading control, mostly the stain-free total protein imaging technology was used as previously described.^30^

### Autophagic flux assays

To assess the effect of PSMI on myocardial autophagic flux, wild type and p62^-/-^ littermate mice at 8 weeks of age were treated with intraperitoneal injections of proteasome inhibitor MG-262 (#A4009, ApexBio; 5 µmol/kg) for 11 hours and then lysosome inhibitor bafilomycin A1 (BFA, 3 µmol/kg, i.p.; #11038, Cayman chemical) for 1 hour or vehicle control (equivalent amount of 60% DMSO in saline). Mice were sacrificed and ventricular myocardial tissue were collected 1 hour after BFA injection for subsequent protein extraction and western blot analyses for LC3, p62, and total ubiquitinated proteins. LC3-II flux and lysosome-mediated total ubiquitinated protein flux were the differential in the myocardial levels of LC3-II or total ubiquitinated proteins between the vehicle control and the BFA-treated subgroups within each genotype and PSMI status group. The LC3-II flux assay in cultured neonatal mouse cardiomyocytes was performed similarly, where lysosome inhibition was achieved with BFA treatment.

### Cytoplasmic and nuclear fractionation of NRVMs

Cytosolic and nuclear proteins were extracted from cultured NRVMs using the Nuclear Extraction Kit (Catalog. #: NC113474; Abcam, Cambridge, MA) according to the manual provided by the manufacturer. In brief, ∼2×10^6^ cells were harvested, washed with PBS, and centrifuged for 5 min at 1000 rpm; the supernatant was discarded. The 200 µl pre-extraction buffer was then added to the cell pellet, mixed, and incubated on ice for 10 min during which the homogenates were vortexed vigorously for 10 sec every 2 min, and then centrifuged at 4°C for 1 min at 12000 rpm. The resultant supernatant was collected as the cytoplasmic protein extract; the pellet was washed with PBS two times, then mixed with 2 volumes of nuclear extraction buffer, and the mixture was incubated on ice for 15 min during which it subject to 5 sec of vortex every 3 min. The extract was further sonicated for 3 × 10 seconds to increase nuclear protein extraction and centrifuged at 14000 rpm for 10 min at 4 °C and the supernatant collected as the nuclear fraction. Both the cytoplasmic and nuclear protein extracts were stored at −80°C until use.

### Immunofluorescence staining and confocal microscopy

Immunofluorescence staining and confocal microscopy were performed on cultured cardiomyocytes or cryosections of mouse myocardium as we previously reported.^14^ In brief, 4% paraformaldehyde-fixed cardiomyocytes on cover glass or myocardial cryosections were washed thrice for 5 min with PBS, incubated with 1% glycine in PBS for at least 30 min, and blocked with 5% BSA at room temperature for 2 h. Primary antibodies were added to specimen and incubated in a moisture chamber at 4 °C overnight; and unbound antibodies were removed via 5 min x 3 washes with PBST before incubation with appropriate secondary antibodies at room temperature for 1 h. The specimens were then rinsed with PBST 5 times for 5 min each. DAPI (Sigma-Aldrich, 1:10) was used for staining nuclei. Then the stained sections were mounted with the VECTASHIELD mounting medium (Vector Laboratories), covered by glass coverslips, sealed with nail polish, and kept in −20°C prior to confocal imaging analyses. The fluorescence staining was visualized and imaged using a confocal microscope (Olympus FluoviewTM 500, Olympus, Waltham, MA).

### RNA isolation, cDNA synthesis and quantitative PCR

Total RNA was isolated as previously described.^15, 22^ The first strand cDNA synthesis was performed using a commercial kit (#4374966, ThemoFisher Scientific, Waltham, MA) as reported.^15, 22^ Transcripts of interest were amplified and quantified using either conventional semi-quantitative PCR (RT-PCR) followed by densitometry or quantitative real-time PCR with the SYBR-Green assay using gene-specific primers purchased from ThemoFisher Scientific (see **Supplementary Table 2** for primer sequences). For conventional RT-PCR, GAPDH was probed as a house-keeping gene control and its level was utilized to normalize the PCR product levels of other genes.^15, 22^ For real-time PCR, data were normalized to mouse acidic ribosomal phosphoprotein P0 (RPLP0) and relative gene expression was calculated using the 2^-ΔΔct^ method.^31^

### Statistical methods

All continuous variables are presented as mean ± SEM unless indicated otherwise. Differences between two groups were evaluated for statistical significance with a 2-tailed unpaired t-test or Welch’s t-test. When the difference among 3 or more groups was evaluated, 1-way ANOVA or, when appropriate, 2-way ANOVA followed by the Tukey test for pairwise comparisons was performed. A p value < 0.05 is considered statistically significant.

## RESULTS

### 1. Increases in myocardial p62, LC3-II, and autophagosomes in Psmc1^CKO^ mice

We have previously reported that pharmacologically induced PSMI activates autophagy in the heart.^13^ Consistent with that, myocardial autophagy is highly activated in mice with early stage proteinopathy which is known to impair UPS function.^14^ However, the impact of cardiac proteasome malfunction on autophagy has not been examined in a genetic model of PSMI. Protease 26S subunit ATPase 1 (PSMC1, also known as rpt2 in yeast and S4 in humans) is one of the six non-redundant AAA-ATPase subunits of the 19S proteasome and is required for the assembly and function of 26S proteasomes.^25, 32^ Using the Cre-Loxp system, we achieved perinatal cardiomyocyte-restricted knockout of the *Psmc1* gene (Psmc1^CKO^) in mice (**Supplementary Figure 1**). In examination of the homozygous Psmc1^CKO^ (αMyHC-Cre::Psmc1^f/f^) mice in comparison with littermate control mice (CTL) consisting of Psmc1^f/+^, Psmc1^f/f^, and heterozygous Psmc1^CKO^ (αMyHC-Cre::PSMC1^f/+^) mice, we confirmed the deletion of the *Psmc1* gene in cardiomyocytes, which resulted in ∼85% reduction of Psmc1 proteins in the heart and the loss of nuclear-enriched Psmc1 staining in cardiomyocytes, but not in non-cardiomyocyte cells (**Supplementary Figure 1B-1D**). Consistent with previous report,^25^ depletion of Psmc1 caused substantial accumulation of myocardial total ubiquitinated proteins and, when coupled with a UPS surrogate substrate GFPdgn,^28^ Psmc1^CKO^ increased the protein level of GFPdgn in heart (**Figure 1A-1C**). We also observed prevalent peri-nuclear p62- and ubiquitin-positive protein aggregates in the cardiomyocytes of Psmc1^CKO^ hearts (**Figure 1D**). Thus, we have created a genetic model of cardiomyocyte-restricted PSMI.

**Figure 1.**
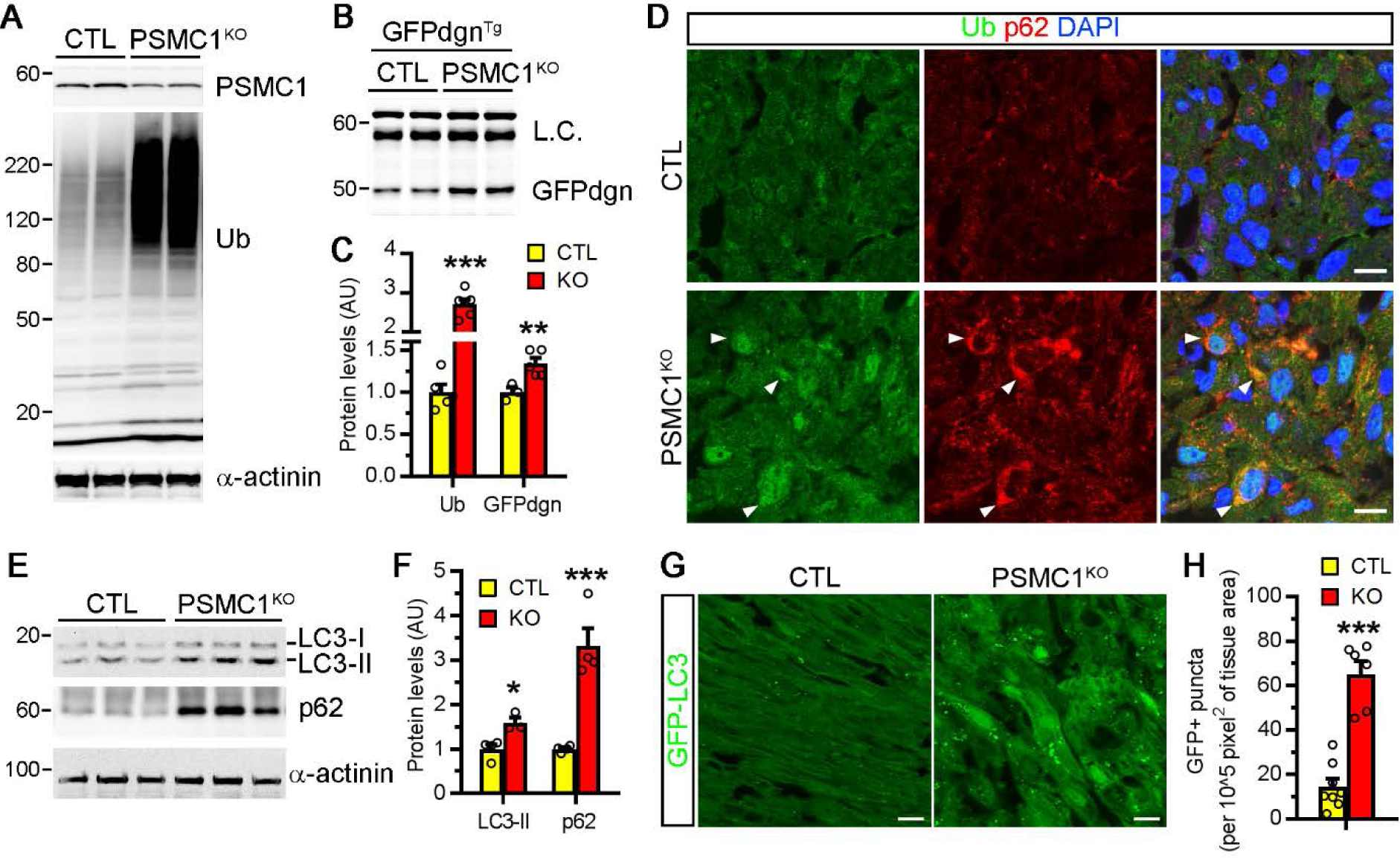
Psmc1^CKO^ increased myocardial p62, LC3-II, and autophagosomes in mice. **A**, Western blot analyses for the indicated proteins in the hearts of CTL and Psmc1^CKO^ mice at postnatal day 2 (P2). Ub, ubiquitin conjugates. **B** and **C**, GFPdgn transgene was introduced into CTL and Psmc1^CKO^ mice. Myocardial GFPdgn protein levels in indicated P2 mice were assessed by western blot (B) and quantified (C). **D**, Immunostaining of ubiquitin (green) and p62 (red) in myocardium sections from P2 mice. Arrowheads mark cardiomyocytes with increased ubiquitinated proteins and p62. Nuclei were counter-stained with DAPI (blue). Bar, 10 µm. **E** and **F**, Western blots (**E**) of indicated proteins in mouse hearts at P2 and the quantification (**F**). **G** and **H**, GFP-LC3 transgene was crossed into CTL and Psmc1^CKO^ mice to label autophagosome (green puncta). Shown are confocal fluorescent images (**G**) of myocardium sections from P2 mice and the quantification of GFP-positive puncta (**H**). Bar, 10 µm. * *p*<0.05, \*\**p*<0.01, \*\*\**p*<0.001 vs CTL. Two-side and unpaired t-test.

Using the Psmc1^CKO^ mice, we then probed the impact of genetic PSMI on the ALP. Loss of Psmc1 led to ∼1.6-fold upregulation of LC3-II and ∼3.3-fold of increase in p62, two key ALP markers, in the heart when compared with the CTL (**Figure 1E, 1F**). p62 is a prototype autophagic receptor that shuttles ubiquitinated proteins to the autophagosome for degradation.^33^ We found the increased p62 frequently co-localized with ubiquitin-positive protein aggregates in Psmc1^CKO^ hearts (**Figure 1D**). To visualize the abundance of autophagosomes, we cross-bred a GFP-LC3 transgene into the control and the Psmc1^CKO^ mice and observed more than 5-fold increase of GFP-positive puncta in Psmc1^CKO^ hearts (**Figure 1G-1H**). Together, these *in vivo* data suggest that genetic PSMI induces the ALP in the heart.

Homozygous Psmc1^CKO^ mice developed severe cardiac failure shortly after birth, as revealed by conscious echocardiography. Morphometric analyses of the echocardiographs recorded at postnatal day 2 showed unchanged left ventricular (LV) end-diastolic posterior wall thickness but increased LV internal diameter at both end-diastole and end-systole, decreased LV ejection fraction (EF), and reduced conscious heart rate in comparison with their littermate controls, including Psmc1^f/+^, Psmc1^f/f^, and heterozygous Psmc1^CKO^ mice. All homozygous Psmc1^CKO^ mice died by postnatal day 10 (**Supplementary Figure 2**). Our data demonstrate that Psmc1 is essential for cardiac UPS function, cardiac development and perinatal survival in mice.

### 2. Upregulation of p62 and autophagic flux in cardiomyocytes by genetic PSMI

The increase of autophagosomes observed in Psmc1^CKO^ hearts could be secondary to cardiac dysfunction and/or a consequence to an impairment in the removal of autophagosomes. To address this question, we silenced *Psmc1* in neonatal rat ventricular myocytes (NRVMs), which led to a remarkable increase in total ubiquitinated proteins and p62, as well as significant depletion of LC3-II (**Figure 2A**). Consistent with our *in vivo* findings, we observed significantly increased cells with GFP-LC3 puncta upon *Psmc1* silencing and more GFP-LC3 puncta in individual *Psmc1*-silenced cells (**Figure 2B, 2C**). To measure autophagic flux, we induced *Psmc1* deletion by infecting neonatal cardiomyocytes isolated from Psmc1^f/f^ mice with adenoviruses expressing *Cre*. LC3-II proteins were significantly reduced in Psmc1-depleted cells, and inhibition of lysosomes with bafilomycin A1 led to a greater increase of LC3-II in these cells when compared with the control cells (**Figure 2D, 2E**), indicative of a remarkable increase of the autophagic flux in cardiomyocytes with genetic PSMI. Combined with our in vivo findings (**Figure 1**), these data indicate that genetic PSMI is sufficient to activate autophagy in cardiomyocytes.

**Figure 2.**
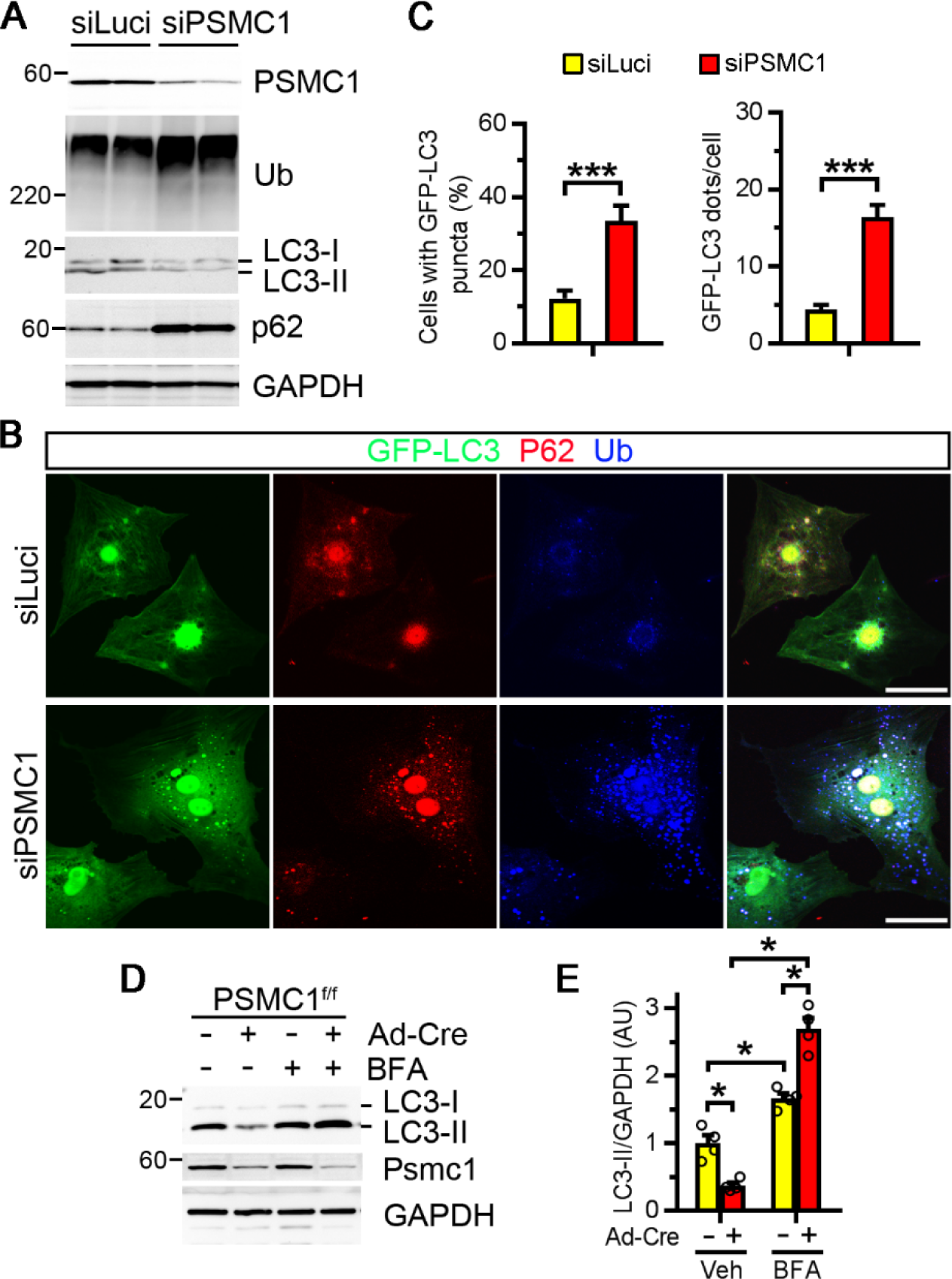
Genetic inhibition of the proteasome increases p62 and autophagic flux in cultured cardiomyocytes. **A-C**, Neonatal rat ventricular myocytes (NRVMs) were transfected with the indicated siRNAs for 72 hours. Cells were infected with Ad-GFP-LC3 for 24 hours before harvest (**B-C**). **A**, Western blot of indicated proteins. **B**, Immunostaining of p62 (red) and Ub (blue) in cells expressing GFP-LC3 (green). **C**, Quantification of cells with GFP-LC3 puncta (%) and the number of puncta in cells expressing GFP-LC3. **D-E**, Neonatal mouse ventricular cardiomyocytes (NMVMs) were isolated from neonatal PSMC1^f/f^ mice and infected with Ad-Gal or Ad-Cre for 72 hours. The cells were treated with vehicle or Bafilomycin A1 (BFA, 100 nM) for 24 hours before harvest. Western blots of indicated proteins (**D**) and the quantification (**E**) are shown. * *P*<0.05, *** *P*<0.001. Two-side and unpaired t-test.

### 3. Activation of myocardial TFEB in mice by both pharmacological and genetic PSMI

The molecular mechanisms underlying the interplay between the UPS and autophagy are incompletely understood. TFEB is a master regulator of autophagy and lysosome function by acting as a transcription factor that induces the expression of a network of genes involved in autophagosome and lysosome biogenesis.^18^ Western blot analysis showed that administration of a bolus of bortezomib (BZM, 10 µg/kg), an FDA-approved proteasome inhibitor, markedly increased myocardial p62 (**Figure 3A, B**) and ubiquitinated proteins in mice (**Supplementary Figure 3**), which was accompanied by a reduction of the slower-migrating TFEBa band and consequently an increase of its faster migrating counterpart (**Figure 3A**), indicative of increases in p62 and the dephosphorylated form of TFEBa by pharmacological PSMI. The dephosphorylated TFEB is prone to nuclear translocation to induce autophagy.^18^ Indeed, subcellular fractionation analyses revealed increases of TFEBa in both cytoplasmic and nuclear fractions of the myocardium from mice treated with BZM, compared with those treated with vehicle control (**Figure 3C ∼ 3E**). In further support of the transactivation of TFEB by PSMI, RT-PCR analyses revealed significantly increases in the mRNA levels of *Uvrag, Vps18, Mcoln1, M6PR* and *p62/Sqstm1*, the well-known target genes of TFEB,^18^ in BZM-treated mouse hearts (**Figure 3F, G**).

**Figure 3.**
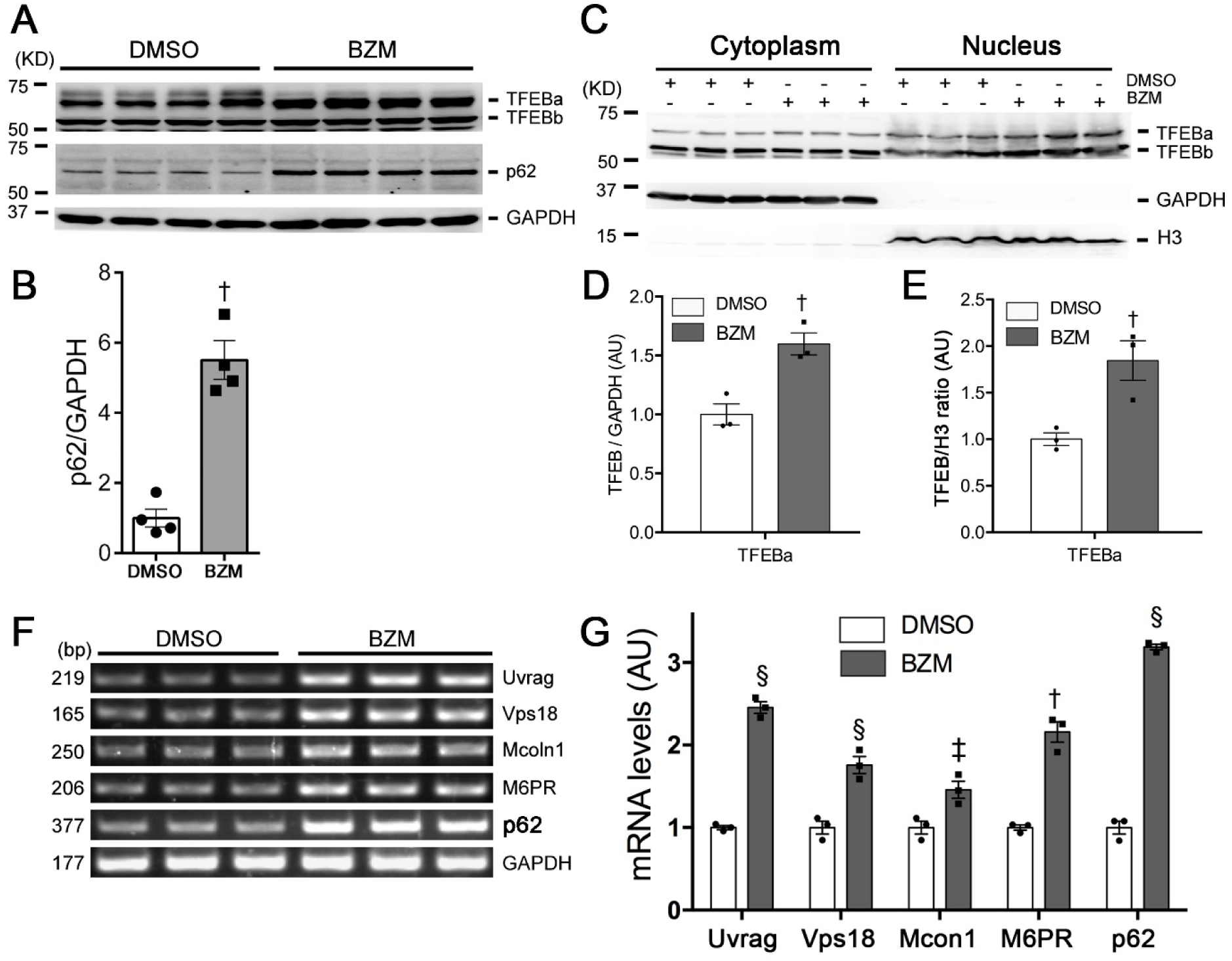
Pharmacological PSMI increases myocardial p62/SQSTM1 and activates TFEB in mice. Mixed sex wild type mice at 5 weeks of age were treated with bortezomib (BZM; 10 µg/kg, i.p.) or vehicle control (60% DMSO in saline). Twelve hours after the treatment, ventricular myocardium was sampled for protein and RNA analyses. **A** and **B**, Representative images (A) of western blot analyses for the indicated proteins and pooled densitometry data of p62 proteins (B). **C-E**, Representative images (C) and densitometry data (D, E) of western blot analyses for TFEB in the cytoplasmic and nuclear fractions of ventricular myocardial proteins. GAPDH and histone H3 (H3) were probed as cytoplasmic and nuclear proteins marks, respectively. **F** and **G**, representative images (F) and pooled densitometry data (G) of RT-PCR analyses for the mRNA levels of indicated genes. Each lane or each dot represents an individual mouse. ^†^p<0.01, ^‡^p<0.001, and ^§^p < 0.0001 vs. the DMSO group, two-sided and unpaired *t*-test.

To determine if genetic PSMI also activates TFEB, we performed immunostaining to analyze TFEB localization in mouse hearts at postnatal day 2 and observed ∼3-fold more cardiomyocytes with nuclear TFEB-positive staining in Psmc1^CKO^ hearts (16.8% of Psmc1^CKO^ vs 5.2% of CTL, *p*<0.01) (**Figure 4A, 4B**). Western blot analysis revealed that depletion of Psmc1 resulted in a significant increase of TFEBa proteins in mouse hearts (**Figure 4C-4D**). Psmc1^CKO^ also significantly increased the transcripts of TFEB target genes including *p62/Sqstm1, MP6R, Uvrag, Vps18* and *Mcoln1* (**Figure 4E**).

**Figure 4.**
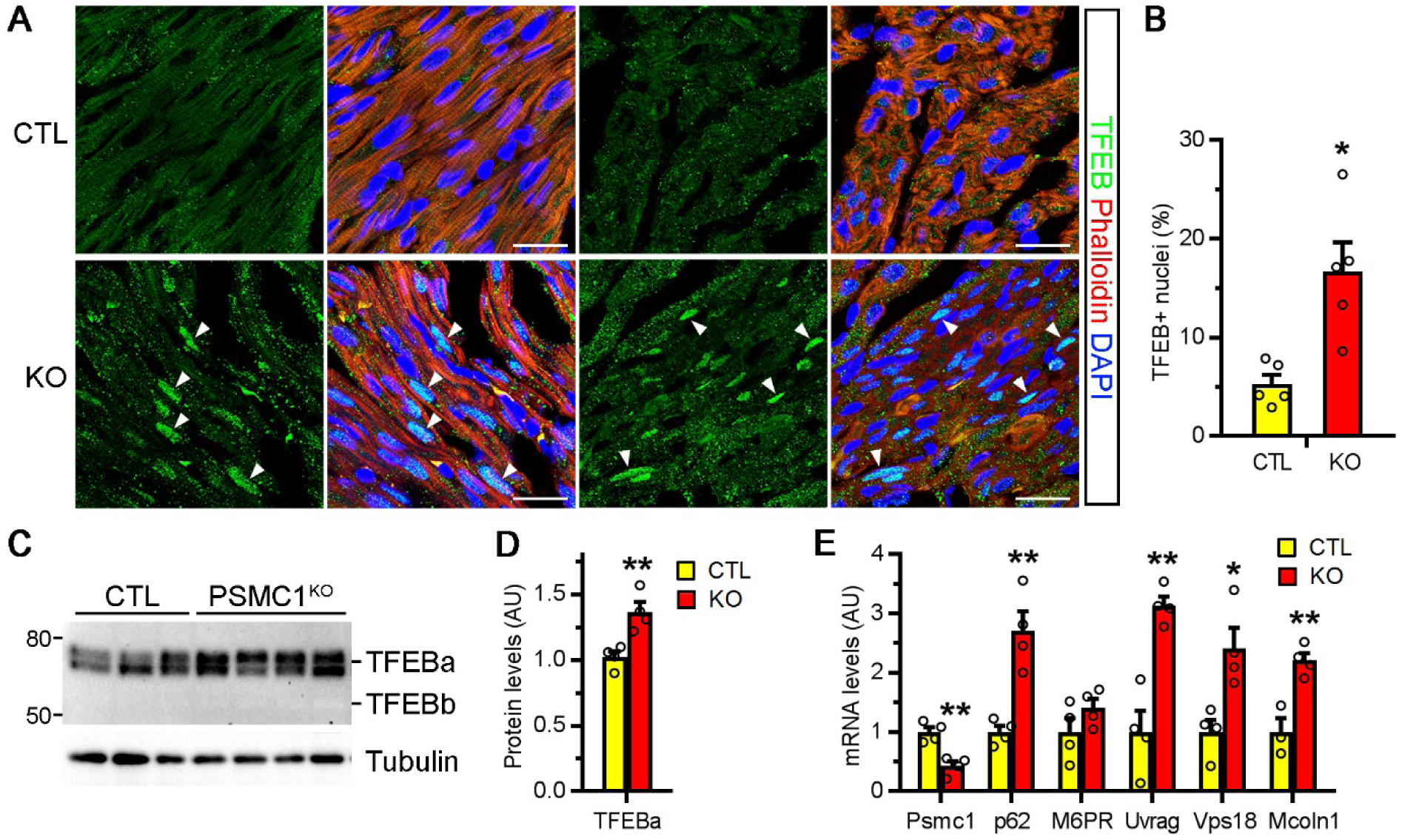
Genetic PSMI via Psmc1^CKO^ activates myocardial TFEB in mice. **A** and **B**, Immunostaining of TFEB (green) in the myocardium sections of CTL and Psmc1^CKO^ mice at P2. Representative confocal micrographs (**A**) and the percentage of TFEB-positive nuclei (arrowheads) were quantified (**B**). The sections were counterstained with Phalloidin (red) and DAPI (blue), respectively. Bar, 20 µm. **C** and **D**, Western blots (**C**) and the quantification (**D**) of TFEB in CTL and Psmc1^CKO^ hearts at P2. **E**, Quantitative real-time PCR analyses of the indicated genes in CTL and Psmc1^CKO^ hearts. * *p*<0.05, ** *p*<0.01 vs. CTL. Two-side and unpaired *t*-test or t-test with Welch correction.

In cultured NRVMs, BZM treatment for 12 and 24 hours also dramatically reduced the slower-migrating TFEBa but increased the faster-migrating TFEBa band (**Figure 5A**), indicative of enhanced dephosphorylation and nuclear translocation. Indeed, subcellular fractionation analyses showed that nuclear TFEBa is the faster-migrating forms and BZM treatment led to a significant accumulation of TFEBa in the nucleus at both 12 and 24 hours (**Figure 5B, C**). Immunostaining for TFEB further confirmed that more BZM-treated cardiomyocytes than the control cells displayed nuclear enrichment of TFEB (**Figure 5D**).

**Figure 5.**
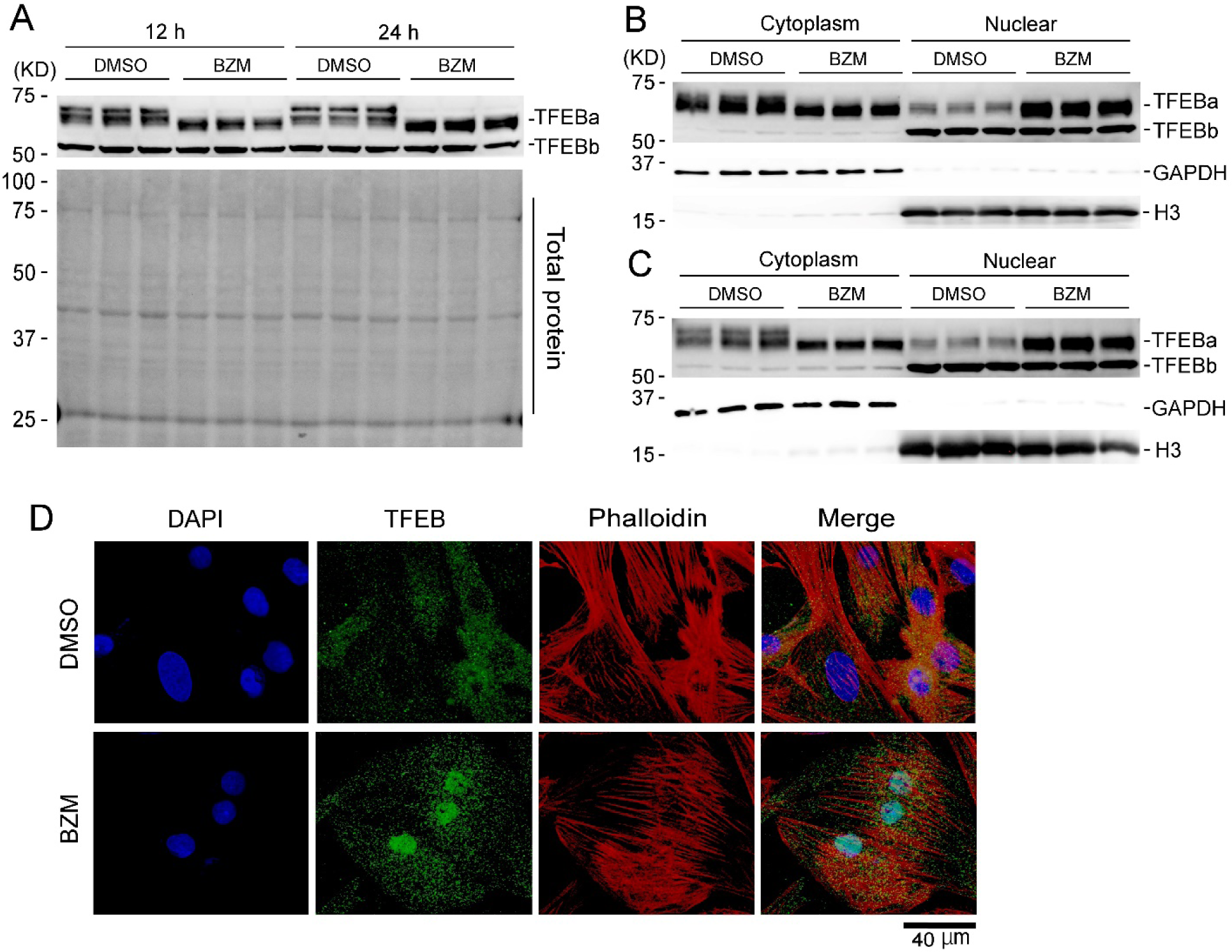
PSMI by BZM leads to dephosphorylation and nuclear translocation of TFEB in cultured neonatal rat ventricular myocytes (NRVMs). Cultured NRVMs treated with BZM (25 nM) or DMSO for 12 or 24 hours were harvested and processed for analyses reported here. **A**, total cellular protein extracts were used for Western blot analyses for TFEB; the stain-free total protein image on the PVDF membrane used for immunoblotting (bottom of A) is used as the loading control. **B** and **C**, cytosolic and nuclear protein extracts from the cultured NRVMs at 12 hours (B) and 24 hours (C) after the treatment were subject to Western blot analyses for the indicated proteins. GAPDH and Histone H3 (H3) were probed as a cytoplasmic and nuclear protein marker, respectively. n= 3 biological repeats/group. **D**, cultured NRVMs grown on cover glasses were fixed with 4% paraformaldehyde at 12 hours after the treatment of DMSO or BZM (25 nM) and used for the indirect immunofluorescence labeling of TFEB (green). The nuclei were stained with DAPI (blue) and F-actin in the cardiomyocytes was stained with Alexa Fluor 568 conjugated phalloidin (red). Representative fluorescence confocal micrographs are shown. Scale bar = 40 μm.

Together, these lines of in vivo and in vitro evidence compellingly demonstrate that both pharmacological and genetic PSMI activate TFEB and its downstream signaling in cardiomyocytes.

### 4. TFEB activation by PSMI in a calcineurin-dependent manner

Phosphorylation of TFEB dictates its subcellular localization and transcriptional activity; the dephosphorylated TFEB translocates from cytoplasm to the nucleus where it initiates downstream gene expression.^18^ We have previously reported that PSMI activates the calcineurin-NFAT pathway in cardiomyocytes and mouse hearts.^34^ Here we observed that the mRNA levels of MCIP1.4, a *bona fide* target gene of the calcineurin-NFAT pathway, were significantly higher in cultured NRVMs and mouse hearts that were treated with a proteasome inhibitor than those in the controls (**Supplementary Figure 4**). Cyclosporine A (CsA) is a potent calcineurin inhibitor. CsA treatment restored the levels of the slower-migrating TFEB bands in the BZM-treated cardiomyocytes in a dose-dependent manner (**Supplementary Figure 5**), indicating that activation of calcineurin is responsible for PSMI-induced TFEB dephosphorylation. Subcellular fractionation (**Figure 6A, B**) and immunostaining (**Supplementary Figure 6)** further confirmed that inhibition of calcineurin by CsA blunted BZM-induced TFEB nuclear translocation. Moreover, CsA treatment attenuated BZM induction of a wide array of TFEB target genes involved in autophagy and lysosome biogenesis, including *Uvrag, Vps18, Beclin 1, Rab7a, p62, cathepsin B, cathepsin D, Lamp1, M6PR* and *Mcoln1* (**Figure 6C, D**). Together, these findings demonstrate that calcineurin activity is required for PSMI to activate TFEB in cardiomyocytes.

**Figure 6.**
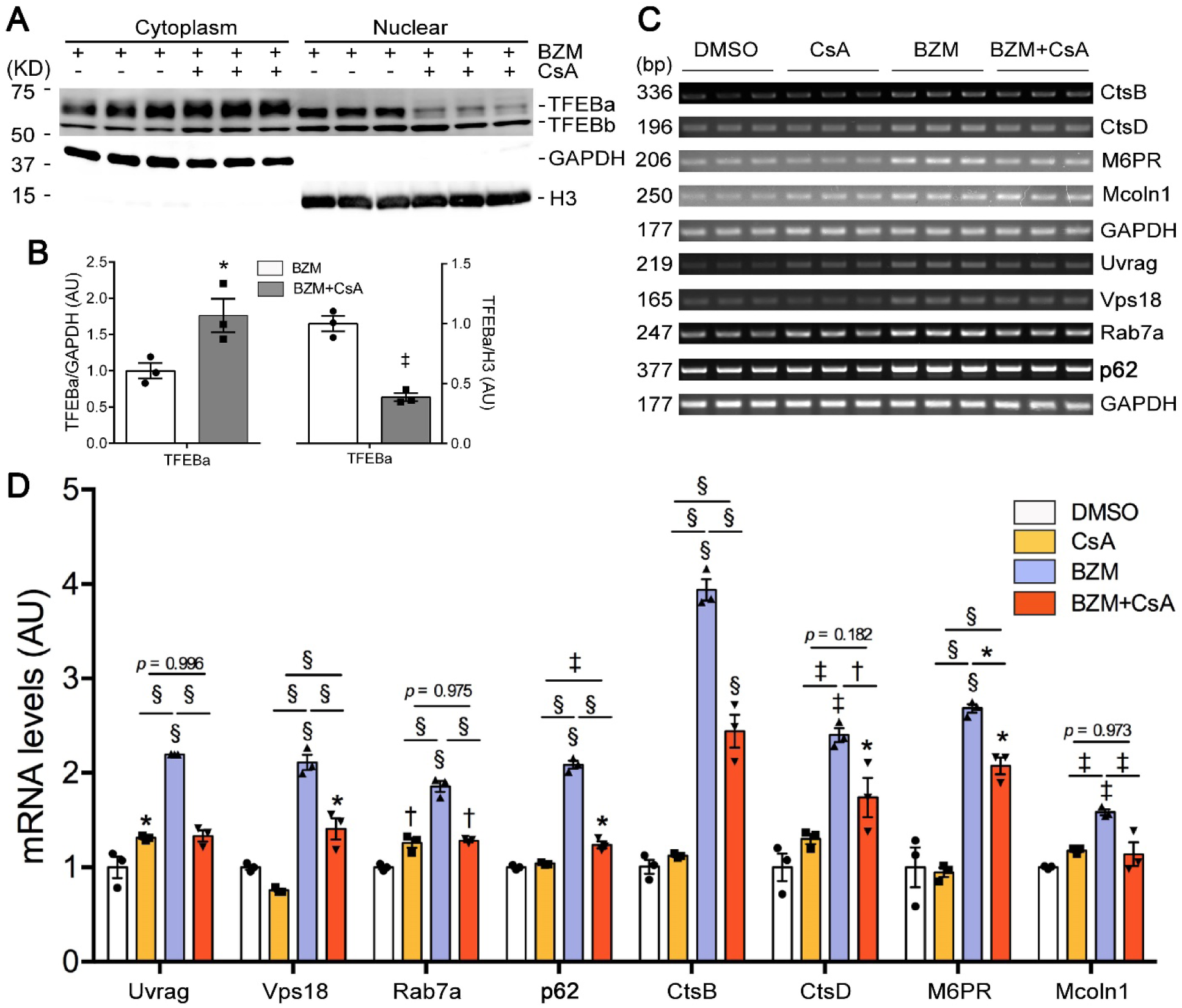
TFEB activation by PSMI in NRVMs is calcineurin-dependent. NRVMs were treated with cyclosporine A (CsA; 1.5 µM) for 4 hours before initiation of the BZM treatment (25 nM). At 12 hours after BZM treatment, cells were harvested for protein or RNA extractions. **A** and **B**, Representative images (A) and densitometry data (B) of western blot analyses for TFEB in the cytosolic and nuclear fractions. GAPDH and histone H3 (H3) were probed as a cytoplasmic and nuclear protein marker, respectively. Welch’s t-test, *p<0.05, ^†^p<0.01, ^‡^p<0.001 vs. the BZM group. **C** and **D**, Representative images (C) and pooled densitometry data (D) of RT-PCR analyses for the mRNA levels of representative TFEB target genes. *p<0.05, ^†^p<0.01, ^‡^p<0.001, and ^§^p < 0.0001 vs. the DMSO group or between the indicated groups; two way ANOVA followed by Tukey’s test. Each dot represents a biological repeat.

### 5. TFEB activation by PSMI requires Mucolipin 1 (Mcoln1)

Mcoln1, also known as transient receptor potential mucolipin 1 (TRPML1), is the main calcium channel on the membrane of late endosomes and lysosomes.^19^ It is purported that during autophagic activation, calcium released from lysosomes via Mcoln1 activates calcineurin, which in turn dephosphorylates TFEB and promotes TFEB nuclear translocation.^19^ To our best knowledge, this has not been demonstrated in cardiomyocytes. We observed significant upregulation of *Mcoln1* gene expression in BZM-treated hearts, Psmc1^CKO^ hearts, and BZM-treated cultured cardiomyocytes (**Figures 3, 4, 6**). We then determined the impact of silencing *Mcoln1* via transfection of *Mcoln1*-specific siRNA on PSMI-induced TFEB activation in cultured NRVMs. Our results showed that silence of *Mcoln1* effectively suppressed BZM-induced TFEB nuclear translocation (**Figure 7A**) and abrogated BZM-induced upregulation of TFEB downstream genes (**Figure 7B, 7C**). Thus, we conclude that PSMI-induced TFEB activation requires Mcoln1.

**Figure 7.**
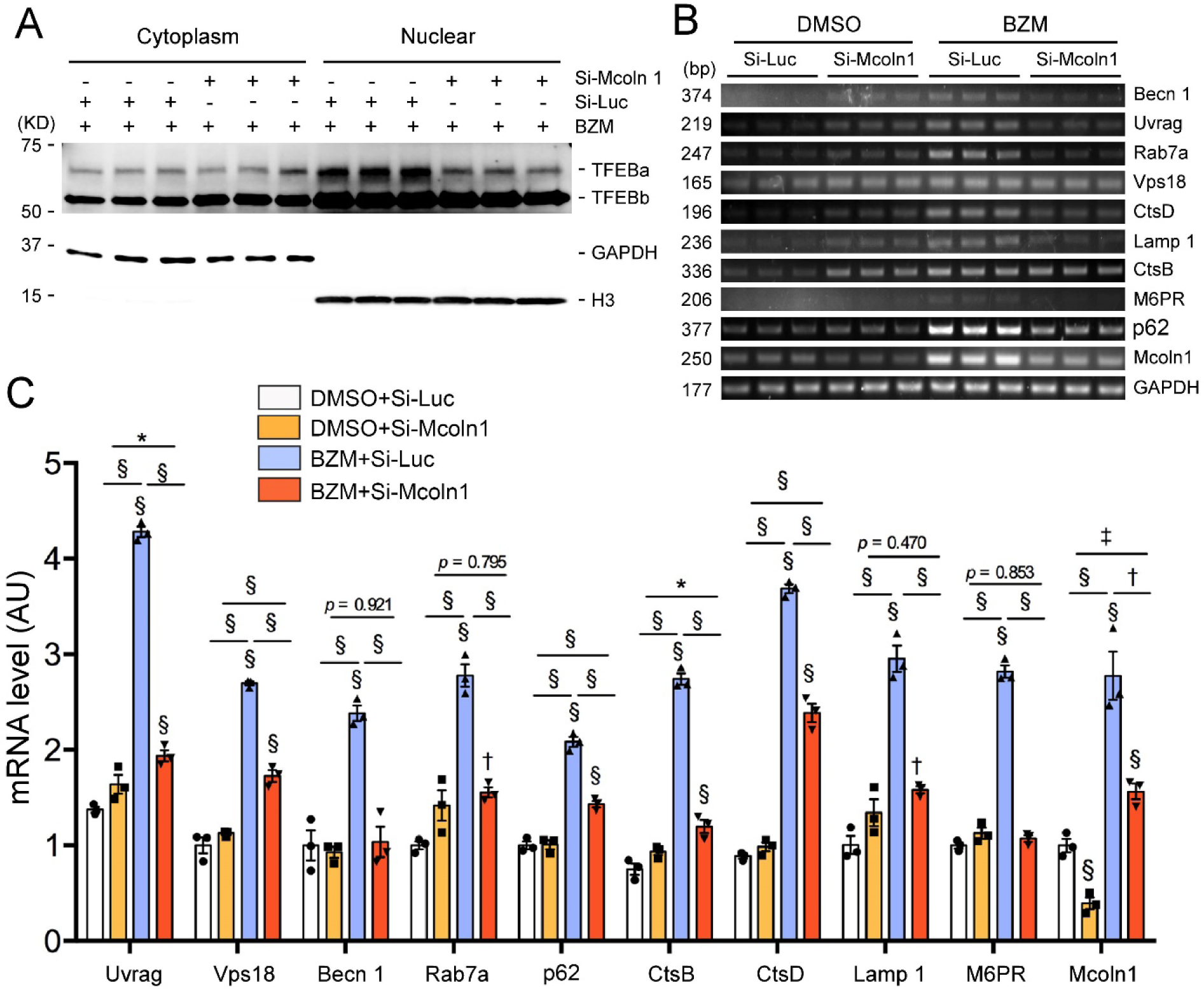
TFEB activation by PSMI in NRVMs is Mcoln1 dependent. At 48 h after transfection of siRNA specific for Mcoln1 (si-Mcoln1) or for luciferase (si-Luc) as the control siRNA, the cells were treated with BZM (25 nM) and harvested 12 h later for cytosolic and nuclear protein fractionation or for total RNA extraction. **A**, Western blot analyses for the indicated proteins. GAPDH and Histone H3 (H3) were probed as a cytoplasmic and nuclear protein marker, respectively. **B** and **C**, representative images (B) and pooled densitometry data (C) of RT-PCR analyses for the mRNA levels of the indicated TFEB target genes. *p<0.05, ^†^p<0.01, ^‡^p<0.001, and ^§^p < 0.0001 vs. the DMSO group or between the indicated groups, n= 3 biological repeats/group; two way ANOVA followed by Tukey tests for pairwise comparisons.

### 6. Requirement of p62 for PSMI to increase autophagy flux in mouse cardiomyocytes and hearts

In general, p62/Sqstm1 is an adaptor protein that abridges ubiquitinated proteins and organelles with autophagosomes for degradation.^33^ PSMI upregulates p62 at both mRNA and protein levels in the heart (**Figures 1E, 2A, 3A, 3B, 3F, 3G and 4E**) but its roles in PSMI-induced autophagy and TFEB activation are not clear. At baseline, myocardial LC3-II flux was comparable between wild type and p62 null mice; PSMI with MG-262 significantly increased LC3-II turnover in wild type mouse hearts but not in p62 null hearts (**Figure 8A∼8C**). To determine whether the effect of p62 ablation observed in mouse hearts is cardiomyocyte-autonomous, we also performed similar tests in cultured mouse cardiomyocytes isolated from wild type and p62 null neonatal mice. Similar to the results from the in vivo tests, PSMI induced a significant increase in LC3-II flux in wild type cardiomyocytes but not in p62 null cardiomyocytes. Different from the in vivo findings, LC3-II flux was significantly lower in p62-null cardiomyocytes than in wild type cardiomyocytes under the basal condition (**Figure 8D∼8F**). Moreover, we also assessed the effects of PSMI with BZM and lysosome inhibition with BFA on the level of steady state ubiquitin conjugates in these cardiomyocytes (**Supplementary Figures 7**). Treatment with either BZM or BFA alone induced a marked increase of total ubiquitinated proteins in both wild type and p62 null cells but the increase in the p62 null cells was much less than in the wild type cells. The treatment combining BZM and BFA showed a discernible additive effect in the wild type but not p62 null cells, indicating that p62 is required for proteasome malfunction to increase the lysosome-mediated clearance of ubiquitinated proteins in cardiomyocytes. Importantly, experiments using cultured NRVMs also revealed that genetic PSMI via siRNA-mediated *Psmc1* silence induced remarkably less accumulation of total ubiquitinated proteins in the cells with siRNA-mediated p62 depletion compared with those without p62 depletion (**Supplementary Figures 8**). Taken together, these in vivo and in vitro experiments demonstrate that compensatory upregulation of autophagic degradation of ubiquitinated proteins under a PSMI or proteasome malfunction state requires p62.

**Figure 8.**
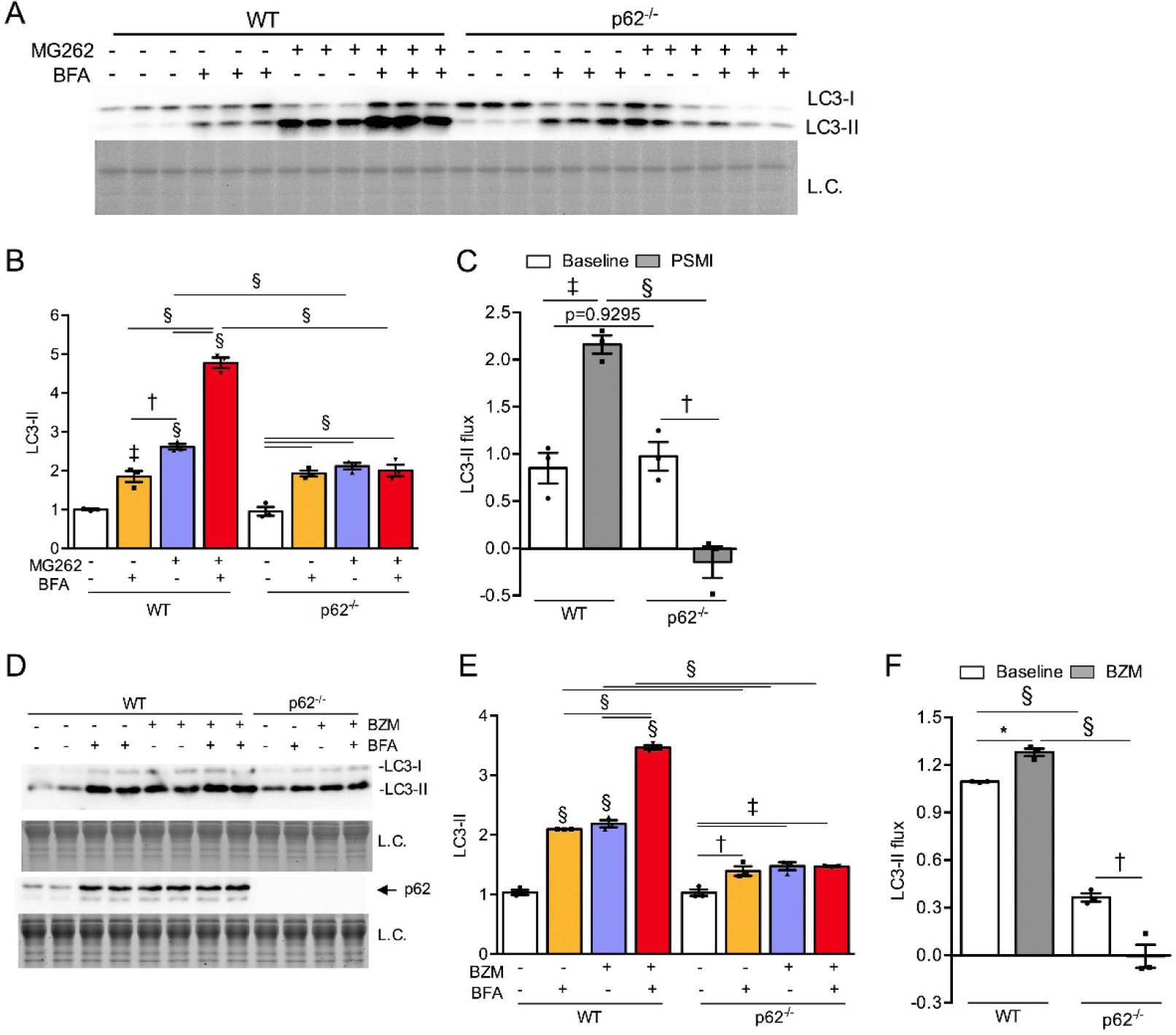
p62/Sqstm1 is required for PSMI to increase autophagic flux in mouse hearts and cardiomyocytes. **A**∼**C**, Wild type (WT) or p62 null (p62^-/-^) mice at 6-8 weeks of age were treated with a proteasome inhibitor MG262 (5 µmol/kg, i.p.) or vehicle control (DMSO); 11 hours later they were treated with bafilomycin A1 (BFA, 3 µmol/kg, i.p.) or vehicle control (DMSO). All the mice were sacrificed for tissue collection at exactly 1 hour after the BFA injection. Shown are the representative images (A) and pooled densitometry data (B) of western blot analyses for LC3. The LC3-II flux derived from the LC-II analyses is presented in panel C. **D**∼**F**, Cardiomyocytes isolated from WT and p62^-/-^ mice at postnatal day 2 were treated with proteasome inhibitor bortezomib (BZM; 25 nM) for 6 hours, followed by bafilomycin A1 (BFA; 25 nM) treatment for additional 6 hours. Total cell lysates were prepared for western blot analyses of LC3, total ubiquitinated proteins and p62 proteins. Loading control (L.C.) used stain-free total protein imaging technology and only a segment of the image is shown in the figure. Data were analyzed by two-way (C) or three-way (B, D) ANOVA followed by Tukey’s post hoc multiple comparisons tests and are presented as Mean ± SEM; n = 3 mice/group. Symbols above the bars indicate the p value for comparison with the vehicle control; \**p*<0.05, †*p*<0.01, ‡*p*<0.001, §*p*<0.0001.

### 7. A positive feedback by p62/Sqstm1 on TFEB activation

p62/Sqstm1 is a TFEB target gene. Indeed, our data presented thus far also compellingly support the notion that induction of p62 by PSMI in cardiomyocytes and mouse hearts is mediated by TFEB activation. Although p62 is just one of the TFEB target genes, PSMI-induced increases in autophagic removal of both LC3-II and ubiquitinated proteins were completely abolished in mouse hearts and cardiomyocytes deficient of p62. This prompted us to speculate that p62 is required for PSMI to sustain TFEB activation; hence, we further compared the induction of TFEB activation by PSMI between wild type and p62 null mice. Immunofluorescence confocal microscopy showed that ablation of p62 significantly attenuated PSMI-induced TFEB nuclear translocation (**Figure 9A**). RT-PCR further showed that induction of representative TFEB target genes in the heart by PSMI was discernibly less effective in p62 null mice than in wild type mice (**Figure 9B, 9C**). These results indicate that p62 as a target gene of TFEB plays a feed-forward role in continuous TFEB activation during PSMI.

**Figure 9.**
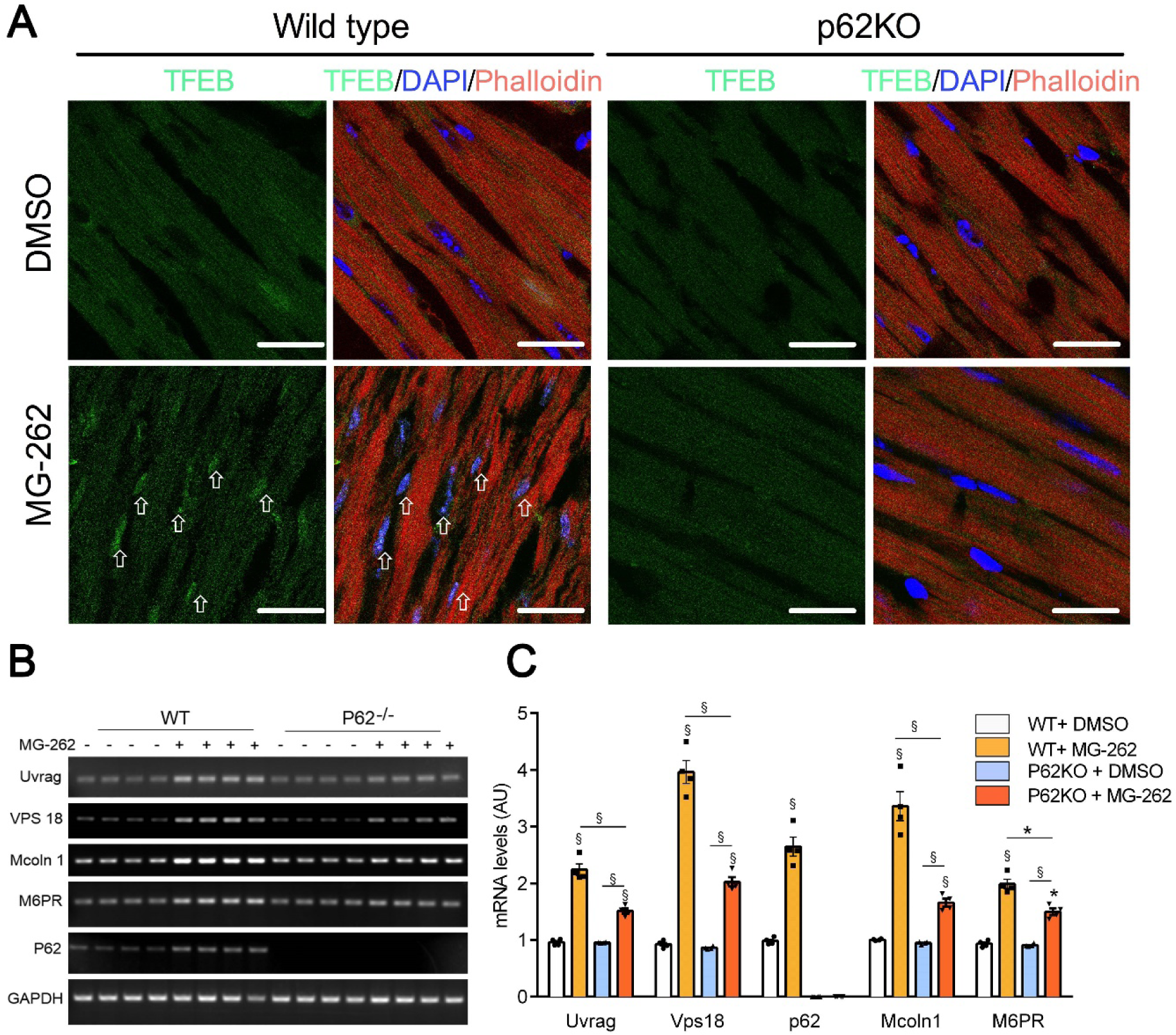
Ablation of p62 (p62KO) attenuates myocardial TFEB activation by PSMI in mice. Wild type and p62KO mice were treated with MG262 or vehicle control as described in Figure 8A before ventricular myocardium was sampled for total RNA extraction or for fixation and further processing to collect cryosections. **A**, Subcellular distribution of TFEB in mouse myocardium. Cryosections from ventricular myocardium were subject to immunofluorescence staining for TFEB (green) as well as counter-staining with DAPI (blue) and Alexa 565-conjugated Phalloidin to reveal the nuclei and F-actin (red), respectively. Shown are representative confocal micrographs. Scale bar = 40µm. **B** and **C**, RT-PCR analyses of representative target genes of TFEB. Total RNA was used for the RT-PCR analyses as described in Figure 3F.

## DISCUSSION

Both the UPS and the ALP are pivotal to protein quality and quantity control in the cell. Targeting these pathways is becoming an attractive strategy for treating human disease, especially for cancer therapies. For example, proteasome inhibitors are highly efficacious in treating multiple myeloma.^35^ However, a significant proportion of patients receiving a proteasome inhibitor-containing regime show cardiotoxicity.^36^ Moreover, UPS and ALP defects are implicated in the pathogenesis of a large subset of heart disease or heart failure.^2^ Therefore, a better understanding of the interplay between these two catabolic pathways is expected to advance cardiac pathophysiology and to identify potentially new therapeutic targets for the related heart disease. Prior studies have suggested that proteasome malfunction increases p62 and activates autophagy in cardiomyocytes but the underlying mechanisms were unclear.^13, 14^ Here we have confirmed this with genetically induced cardiac PSMI. More importantly, we have delineated for the first time the Mcoln1-calcineurin-TFEB-p62 pathway in mediating the autophagic activation by proteasome malfunction and discovered that Mcoln1 and p62 play an essential feed-forward role in TFEB activation by PSMI. Activation of autophagy apparently acts to minimize the toxicity resulting from proteasome malfunction and thereby helps maintain proteostasis in the cell. These discoveries provide significant mechanistic insight into cardiac proteostasis.

### 1. Proteasome malfunction activates the ALP

Recent studies suggest the existence of a coordinated and complementary crosstalk between the UPS and the ALP. Inhibition of proteasome using either pharmacological compounds or genetic means resulted in the upregulation of the autophagic activity in various types of cells in vitro,^37, 38^ including cardiomyocytes.^13, 39, 40^ In vivo, inactivation of proteasome via expression of a dominant negative mutant of β2 proteasome subunit in Drosophila eyes led to tissue-restricted autophagy activation in HDAC6-dependent manner.^41^ In mammals, both proteasome malfunction and activation of autophagy are observed in the heart of well-established mouse models of cardiac proteinopathy and a mouse model of hypertrophic cardiomyopathy that carries a mutant of myosin-binding protein C.^14, 42^ Supporting a functional link between the UPS and the ALP in myocardium, administration of BZM to mice increased autophagosomes and autophagic flux in the heart.^13^ Although directly targeting immunoproteasome subunits (β2i and β5i) had been attempted in mice, such modifications do not alter global proteasomal proteolytic activities.^43, 44^ Thus far, in vivo genetic evidence that directly supports ALP activation by proteasome malfunction in the heart is still lacking. This critical gap is now filled by the present study.

An important contribution of this study is to have achieved and characterized cardiomyocyte-restricted ablation of an essential proteasome subunit gene (*Psmc1*) for the first time in vertebrates (**Supplementary Figure 1, 2**), which confirms with a genetic approach that PSMI activates autophagy (**Figure 1** and **2**). As one of the 19S proteasome subunits, PSMC1 is needed for the formation and activation of 26S proteasomes. Deletion of *Psmc1* in neurons caused neuronal degeneration in mice.^25^ Here we report that targeting *Psmc1* in mouse hearts also led to severe proteasome impairment, as evidenced by the drastic accumulation of ubiquitinated proteins and protein aggregates and by a significant increase of the proteasome surrogate substrate (GFPdgn) in Psmc1-deficient hearts. Upon proteasome inactivation, we observed a remarkable increase of autophagosomes and an upregulation of autophagic proteins, including p62 and LC3-II in the heart. Moreover, p62 frequently co-localized with ubiquitinated proteins and autophagosomes in Psmc1-deficient cardiomyocytes, representing an intermediate state of these cargos *en route* to lysosomal degradation. These lines of evidence, coupled with the increased autophagic flux in Psmc1-deficient cardiomyocytes, demonstrate that autophagy is activated by genetically induced proteasome malfunction in the heart. Apparently, the ALP activation is not sufficient to compensate loss of proteasome function in the heart because perinatal *Psmc1* ablation in cardiomyocytes resulted in severe dilated cardiomyopathy and mouse premature death (**Supplementary Figure 2**).

### 2. Proteasome malfunction activates TFEB

The mechanistic link between the UPS and autophagy remains incompletely delineated. One of the most important contributions of this work is the discovery that PSMI activates TFEB, the master transcription regulator for the ALP. First, we found that pharmacological PSMI increases myocardial TFEB protein levels, TFEB nuclear translocation, and TFEB target gene expression (**Figure 3**); second, genetic PSMI via Psmc1^CKO^ increased TFEB nuclear localization and TFEB target gene expression in mice (**Figure 4**); and lastly, PSMI induced rapidly TFEB dephosphorylation, nuclear translocation, target gene expression in a cardiomyocyte-autologous manner (**Figures 5, 6**). Thus, both in vitro and in vivo evidence compellingly demonstrate a rapid activation of TFEB by proteasome malfunction.

A study using fruit flies showed that HDAC6 mediated autophagic activation by proteasome inhibition;^41^ however, PSMI does not appear to upregulate HDAC6 in mammalian cells.^45^ Several transcription factors such as p53,^46, 47^ Nrf2,^48^ and NFκB,^49, 50^ have been shown to regulate the transcription of a subset of autophagy genes. Proteasome malfunction may lead to accumulation of these transcription factors, which may in turn play a role in autophagy activation. Nevertheless, it remains unclear whether upregulation or activation of a few of these autophagic proteins is sufficient to facilitate autophagic degradation of the proteins that were originally deemed for proteasomal degradation. The activation of TFEB is capable of upregulating coordinately the full spectrum of ALP genes required for sustained autophagic degradation. Hence, our identification of a previously unrecognized role of TFEB in mediating the crosstalk between the UPS and ALP is highly significant because increasing TFEB has been shown to confer cardioprotection under a number of pathological conditions.^21, 22, 51^

### 3. Proteasome malfunction activates TFEB via Mcoln1-Calcineurin

In TFEB activation by starvation, calcineurin which is activated by the calcium released from the lysosome via Mcoln1, is responsible for the dephosphorylation of TFEB.^19^ The role of calcineurin in the activation of TFEB by PSMI has not been examined in any cell types before. We previously showed the activation of the calcineurin-NFAT pathway in cultured cardiomyocytes by PSMI and in the heart of mice with UPS functional insufficiency.^34^ And this is further confirmed in the present study because the mRNA levels of MCIP1.4/Rcan1, a *bona fide* target gene of the calcineurin-NFAT pathway, were significantly increased by PSMI in both cultured cardiomyocytes and intact mice (**Supplementary Figure 4**). This increased calcineurin activity plays an important role in mediating the TFEB activation by PSMI because inhibition of calcineurin phosphatase activity with CsA diminished PSMI-induced TFEBa dephosphorylation in a dose-dependent fashion (**Supplementary Figure 5**) and markedly reduced PSMI-induced TFEB nuclear localization and target gene expression (**Figure 6**).

The role of Mcoln1 in TFEB activation was not investigated previously in cardiomyocytes; moreover, the role of Mcoln1 in the activation of TFEB by PSMI has not been examined in any cell types before. Here we found that as a target gene of TFEB, Mcoln1 expression was significantly increased by PSMI whereas prevention of this increase in Mcoln1 with specific siRNA abolished PSMI-induced TFEBa nuclear translocation and the expression of TFEB target genes in cultured cardiomyocytes (**Figure 7**), demonstrating for the first time that Mcoln1 is indispensable in the TFEB activation by PSMI. This also indicates that calcium release from lysosomes to the cytosol via Mcoln1 plays an essential role in the activation of the phosphatase calcineurin by PSMI and that Mcoln1 exerts a feedforward role in the TFEB activation by PSMI. Since it has been reported that calcineurin is degraded by the UPS and calcineurin proteins accumulate in mouse hearts with UPS impairment,^34, 52^ it is highly likely that PSMI increases calcineurin activities through both stabilization of calcineurin proteins and, via Mcoln1, facilitation of calcineurin activation. As a lysosomal calcium channel, Mcoln1 can be activated by lysosomal stress.^53^ Upon PSMI, the share of ubiquitinated proteins that goes to the ALP for degradation is dramatically increased and thereby imposes stress on lysosomes. Mcoln1 can act as a reactive oxygen species (ROS) sensor and activated by increased ROS;^54^ hence, the activation of Mcoln1 by PSMI may also be through increasing ROS as PSMI has been shown to increase ROS production in the cell.^55^

### 4. The role of p62/Sqstm1 in the activation of autophagy by PSMI

We previously reported upregulation of myocardial p62 and autophagy in mouse models of cardiac proteinopathy induced by cardiomyocyte-restricted overexpression of human disease linked mutant desmin or CryAB^R120G^.^14^ Here we collected both in vivo and in vitro evidence that both pharmacological and genetic PSMI upregulate both mRNA and protein levels of p62 in cardiomyocytes (**Figures 1∼3, 6, 8**). At least a part of this upregulation is mediated by TFEB because inhibition of TFEB activation via either calcineurin inhibition or Mcoln1 knockdown blunted the induction of p62 expression by PSMI (**Figures 6, 7**). A recent study by Sha *et al*. detected in SH-SY5Y neuroblastoma cells and RPMI 8226 myeloma cells that a short exposure to proteasome inhibitor BZM (10 nM/13h or 1 µM/4h) could lead to a rapid increase in the mRNA levels of *p62* and *GABARAPL1* but not in other autophagy related genes or lysosomal genes although longer exposure (20h) to BZM (100nM) could induce most of these genes.^45^ This is in agreement with our findings that marked upregulation of both p62 and other ALP related genes was detected in cultured NRVMs that have been treated with BZM (25 nM) for 12h. In stark contrast to our findings in cardiomyocytes, their further experiments showed siRNA-mediated TFEB knockdown did not alter the induction of p62 by BZM treatment in M17 cells,^45^ suggesting the mechanisms by which PSMI upregulates p62 may be cell-type specific.

On one hand, p62 can bind ubiquitinated substrates via its ubiquitin-associated domain (UBA) and condensate the bound cargos through its PB1-domain-mediated self-oligomerization;^3^ on the other hand, p62 recruits and activates the core autophagosome machinery (e.g., the ULK1-ATG1 complex) via interacting with the claw domain of FIP200 and recruits ATG8/LC3 decorated phagophore via its LC3-interacting region (LIR).^33, 56^ Hence, p62 acting as the prototype receptor for ubiquitinated cargos plays likely an important role in cargo-initiated selective autophagy. However, the in vivo necessity of p62 in cardiac protein quality control at baseline and during proteotoxic stress has not been established until the present study. Here we detected that the basal level of myocardial lysosome-mediated LC3-II flux was comparable between wild type and p62 null mice but the responses of the LC3-II flux to PSMI were drastically different between wild type and p62 null mice (**Figure 8A∼8C**). During PSMI, myocardial LC3-II flux was substantially increased in wild type mice but the flux became completely undetectable in the p62 null mice. Similar findings were obtained in cultured wild type and p62 null neonatal mouse cardiomyocytes but, notably, p62-deifciency significantly decreased LC3-II flux and the lysosome-mediated ubiquitinated proteins flux under both the control culturing condition and during PSMI (**Figure 8D∼8F, Supplementary Figure 7**). This difference between in vivo and cell culture studies is likely caused by that cardiomyocytes when cultured in vitro experience inevitable stresses even in absence of exposure to a proteasome inhibitor. Taken together, these new in vivo and in vitro findings support compellingly that p62 is required for proteasome malfunction to increase autophagy and for lysosomal removal of ubiquitinated proteins in cardiomyocytes and hearts during proteasome malfunction.

There is strong evidence that the rapid upregulation of p62 by PSMI helps channel the ubiquitinated proteins to the ALP for degradation and thereby alleviates the accumulation of ubiquitinated proteins resulting from proteasome malfunction. p62 does so through promoting the aggregation of ubiquitinated proteins and targeting the ubiquitinated cargoes to the ALP. Besides binding to ubiquitinated cargos and targeting them selectively to the ALP, p62 was recently shown to recognize and bind the N-degron in proteins that are normally degraded by the UPS via the N-end rule pathway.^57^ Hence, upon proteasome impairment, the accumulated proteins with an N-degron may use the upregulated p62 as the bridge to the ALP pathway in a ubiquitin-independent manner. Nevertheless, the rapid upregulation of p62 by PSMI requires ubiquitination, according to a recent report.^45^ Consistent with findings from prior studies using cultured cells,^45^ including cultured cardiomyocytes,^14^ our in vivo and in vitro work revealed that PSMI-induced increases in the steady state total ubiquitinated proteins in hearts (*data not shown*) and cultured cardiomyocytes (**Supplementary Figures 7**) were significantly less in p62 null mice compared with wild type mice and this was recapitulated in NRVMs subjected to siRNA-mediated *Psmc1* and *p62* silence (**Supplementary Figure 8**). Taken together with the lysosome-mediated ubiquitinated proteins flux data, these findings indicate that significantly more ubiquitinated proteins are segregated/stabilized by p62 than those removed by p62-triggered autophagy in cardiomyocytes with proteasome malfunction.

### 5. The role of p62/Sqstm1 in the activation of TFEB by PSMI

An unexpected finding of this study is that p62 positively feeds back on TFEB activation in the heart, as evidenced by that cardiac TFEB nuclear localization and target gene expression induced by PSMI were remarkably attenuated in the p62 null mice than the WT mice (**Figure 9**). The mechanism by which p62 does so is currently unclear but p62 is a known singling hub and can interact with a large number of signaling proteins.^58^ To this end, it is worthy to note that p62 as an amino acid sensor can be recruited to and then helps activate mTOR complex 1 (mTORC1);^59^ hence, the positive regulation of TFEB activity by p62 should not be mediated by a mTORC1-centered pathway because activation of mTORC1 inactivates TFEB.^18^ The consumption of lysosomal machinery by p62-mediated increases of autophagy during PSMI can certainly activate TFEB to replenish lysosomes.

### 6. Conclusions

In summary, the present study delineates a molecular pathway for proteasome malfunction to activate autophagy in cardiomyocytes (**Figure 10**), which posts that proteasome malfunction induces calcineurin-mediated activation of TFEB and thereby increases p62 and the selective removal of ubiquitinated proteins by autophagy. We have also presented the first definitive *in vivo* evidence that p62 is required for proteasome malfunction to activate autophagy and the upregulation of p62 by proteasome malfunction exerts a feedforward effect on TFEB activation in the heart.

**Figure 10.**
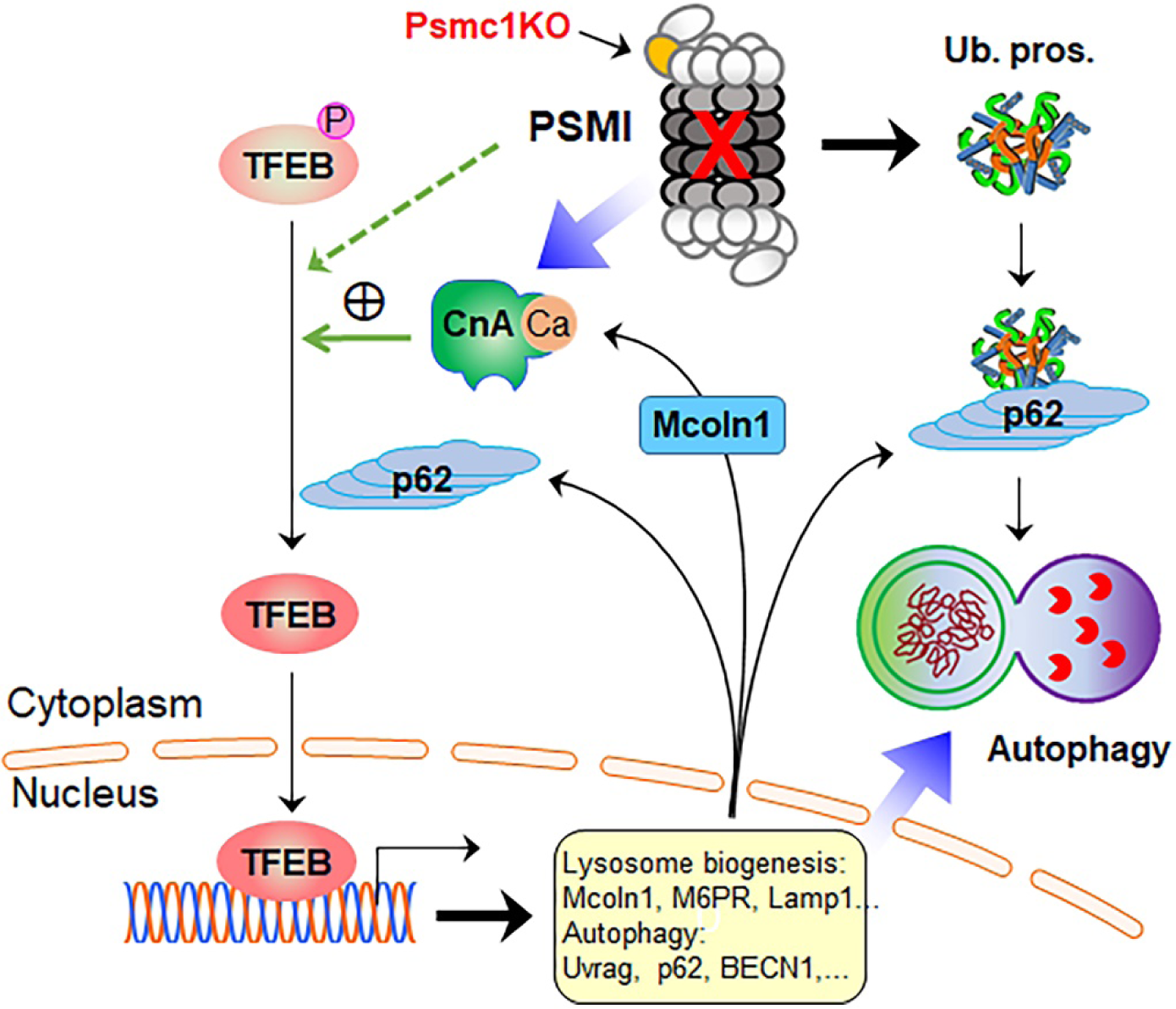
An overall model for proteasome malfunction to activate autophagy through the Mcoln1-calcineurin-TFEB-p62 pathway. This model posts that proteasome malfunction accumulates and activates calcineurin, which in turn dephosphorylates and activates TFEB; the activation of TFEB increases the expression of Mcoln1, p62 and other genes of the CLEAR network and thereby increases autophagy. Mcoln1 and p62 in turn exert a feed-forward effect on TFEB activation whereas p62 recruits and condenses ubiquitinated proteins for autophagic degradation. PSMI, proteasome inhibition; CnA, calcineurin; Psmc1KO, cardiomyocyte-restricted knockout of *Psmc1*; Ub. pro., ubiquitinated proteins.

## Non-standard Abbreviations and Acronyms

ALP: autophagic-lysosomal pathway
BFA: bafilomycin A1
CLEAR: the coordinated lysosomal expression and regulation element
CTL: control
KO: knock out
mTORC1: mechanistic target of rapamycin complex 1
Psmc1-cKO: cardiomyocyte-restricted knockout of the *Psmc1* gene
PSMI: proteasome inhibition
UPS: ubiquitin-proteasome system

## Acknowledgements

This study is in part supported by NIH R01 grants HL072166, HL085629 (to X.W.), and HL131667 (to X.W., T.C.), as well as HL124248 (to H.S.).

## References

1. Wang X, Pattison JS and Su H. Posttranslational modification and quality control. Circ Res. 2013;112:367–81.

2. Wang X and Robbins J. Proteasomal and lysosomal protein degradation and heart disease. J Mol Cell Cardiol. 2014;71:16–24.

3. Wang C and Wang X. The interplay between autophagy and the ubiquitin-proteasome system in cardiac proteotoxicity. Biochim Biophys Acta. 2015;1852:188–94.

4. Collins GA and Goldberg AL. The Logic of the 26S Proteasome. Cell. 2017;169:792–806.

5. Day SM. The ubiquitin proteasome system in human cardiomyopathies and heart failure. Am J Physiol Heart Circ Physiol. 2013;304:H1283–93.

6. Weekes J, Morrison K, Mullen A, Wait R, Barton P and Dunn MJ. Hyperubiquitination of proteins in dilated cardiomyopathy. Proteomics. 2003;3:208–16.

7. Li J, Horak KM, Su H, Sanbe A, Robbins J and Wang X. Enhancement of proteasomal function protects against cardiac proteinopathy and ischemia/reperfusion injury in mice. J Clin Invest. 2011;121:3689–700.

8. Tian Z, Zheng H, Li J, Li Y, Su H and Wang X. Genetically induced moderate inhibition of the proteasome in cardiomyocytes exacerbates myocardial ischemia-reperfusion injury in mice. Circ Res. 2012;111:532–42.

9. Li J, Powell SR and Wang X. Enhancement of proteasome function by PA28α overexpression protects against oxidative stress. FASEB J. 2011;25:883–93.

10. Rajagopalan V, Zhao M, Reddy S, Fajardo GA, Wang X, Dewey S, Gomes AV and Bernstein D. Altered Ubiquitin-Proteasome Signaling in Right Ventricular Hypertrophy and Failure. Am J Physiol Heart Circ Physiol. 2013;305:H551–62.

11. Ranek MJ, Zheng H, Huang W, Kumarapeli AR, Li J, Liu J and Wang X. Genetically induced moderate inhibition of 20S proteasomes in cardiomyocytes facilitates heart failure in mice during systolic overload. J Mol Cell Cardiol. 2015;85:273–81.

12. Li J, Ma W, Yue G, Tang Y, Kim IM, Weintraub NL, Wang X and Su H. Cardiac proteasome functional insufficiency plays a pathogenic role in diabetic cardiomyopathy. J Mol Cell Cardiol. 2017;102:53–60.

13. Zheng Q, Su H, Tian Z and Wang X. Proteasome malfunction activates macroautophagy in the heart. Am J Cardiovasc Dis. 2011;1:214–26.

14. Zheng Q, Su H, Ranek MJ and Wang X. Autophagy and p62 in cardiac proteinopathy. Circ Res. 2011;109:296–308.

15. Pan B, Lewno MT, Wu P and Wang X. Highly Dynamic Changes in the Activity and Regulation of Macroautophagy in Hearts Subjected to Increased Proteotoxic Stress. Front Physiol. 2019;10:758.

16. Liu J, Chen Q, Huang W, Horak KM, Zheng H, Mestril R and Wang X. Impairment of the ubiquitin-proteasome system in desminopathy mouse hearts. FASEB J. 2006;20:362–4.

17. Chen Q, Liu JB, Horak KM, Zheng H, Kumarapeli AR, Li J, Li F, Gerdes AM, Wawrousek EF and Wang X. Intrasarcoplasmic amyloidosis impairs proteolytic function of proteasomes in cardiomyocytes by compromising substrate uptake. Circ Res. 2005;97:1018–26.

18. Wang X and Cui T. Autophagy modulation: a potential therapeutic approach in cardiac hypertrophy. Am J Physiol Heart Circ Physiol. 2017;313:H304–H319.

19. Medina DL, Di Paola S, Peluso I, Armani A, De Stefani D, Venditti R, Montefusco S, Scotto-Rosato A, Prezioso C, Forrester A, Settembre C, Wang W, Gao Q, Xu H, Sandri M, Rizzuto R, De Matteis MA and Ballabio A. Lysosomal calcium signalling regulates autophagy through calcineurin and TFEB. Nat Cell Biol. 2015;17:288–99.

20. Palmieri M, Impey S, Kang H, di Ronza A, Pelz C, Sardiello M and Ballabio A. Characterization of the CLEAR network reveals an integrated control of cellular clearance pathways. Hum Mol Genet. 2011;20:3852–66.

21. Ma X, Mani K, Liu H, Kovacs A, Murphy JT, Foroughi L, French BA, Weinheimer CJ, Kraja A, Benjamin IJ, Hill JA, Javaheri A and Diwan A. Transcription Factor EB Activation Rescues Advanced alphaB-Crystallin Mutation-Induced Cardiomyopathy by Normalizing Desmin Localization. J Am Heart Assoc. 2019;8:e010866.

22. Pan B, Zhang H, Cui T and Wang X. TFEB activation protects against cardiac proteotoxicity via increasing autophagic flux. J Mol Cell Cardiol. 2017;113:51–62.

23. Ma X, Godar RJ, Liu H and Diwan A. Enhancing lysosome biogenesis attenuates BNIP3-induced cardiomyocyte death. Autophagy. 2012;8:297–309.

24. Godar RJ, Ma X, Liu H, Murphy JT, Weinheimer CJ, Kovacs A, Crosby SD, Saftig P and Diwan A. Repetitive stimulation of autophagy-lysosome machinery by intermittent fasting preconditions the myocardium to ischemia-reperfusion injury. Autophagy. 2015;11:1537–60.

25. Bedford L, Hay D, Devoy A, Paine S, Powe DG, Seth R, Gray T, Topham I, Fone K, Rezvani N, Mee M, Soane T, Layfield R, Sheppard PW, Ebendal T, Usoskin D, Lowe J and Mayer RJ. Depletion of 26S proteasomes in mouse brain neurons causes neurodegeneration and Lewy-like inclusions resembling human pale bodies. J Neurosci. 2008;28:8189–98.

26. Agah R, Frenkel PA, French BA, Michael LH, Overbeek PA and Schneider MD. Gene recombination in postmitotic cells. Targeted expression of Cre recombinase provokes cardiac-restricted, site-specific rearrangement in adult ventricular muscle in vivo. J Clin Invest. 1997;100:169–79.

27. Komatsu M, Waguri S, Koike M, Sou YS, Ueno T, Hara T, Mizushima N, Iwata J, Ezaki J, Murata S, Hamazaki J, Nishito Y, Iemura S, Natsume T, Yanagawa T, Uwayama J, Warabi E, Yoshida H, Ishii T, Kobayashi A, Yamamoto M, Yue Z, Uchiyama Y, Kominami E and Tanaka K. Homeostatic levels of p62 control cytoplasmic inclusion body formation in autophagy-deficient mice. Cell. 2007;131:1149–63.

28. Kumarapeli AR, Horak KM, Glasford JW, Li J, Chen Q, Liu J, Zheng H and Wang X. A novel transgenic mouse model reveals deregulation of the ubiquitin-proteasome system in the heart by doxorubicin. FASEB J. 2005;19:2051–3.

29. Mizushima N, Yamamoto A, Matsui M, Yoshimori T and Ohsumi Y. In vivo analysis of autophagy in response to nutrient starvation using transgenic mice expressing a fluorescent autophagosome marker. Mol Biol Cell. 2004;15:1101–11.

30. Zhang H, Pan B, Wu P, Parajuli N, Rekhter MD, Goldberg AL and Wang X. PDE1 inhibition facilitates proteasomal degradation of misfolded proteins and protects against cardiac proteinopathy Sci Adv. 2019:(in press).

31. Zou J, Ma W, Li J, Littlejohn R, Zhou H, Kim IM, Fulton DJR, Chen W, Weintraub NL, Zhou J and Su H. Neddylation mediates ventricular chamber maturation through repression of Hippo signaling. Proc Natl Acad Sci U S A. 2018;115:E4101–E4110.

32. Glickman MH and Ciechanover A. The ubiquitin-proteasome proteolytic pathway: destruction for the sake of construction. Physiol Rev. 2002;82:373–428.

33. Turco E, Witt M, Abert C, Bock-Bierbaum T, Su MY, Trapannone R, Sztacho M, Danieli A, Shi X, Zaffagnini G, Gamper A, Schuschnig M, Fracchiolla D, Bernklau D, Romanov J, Hartl M, Hurley JH, Daumke O and Martens S. FIP200 Claw Domain Binding to p62 Promotes Autophagosome Formation at Ubiquitin Condensates. Mol Cell. 2019;74:330–346 e11.

34. Tang M, Li J, Huang W, Su H, Liang Q, Tian Z, Horak KM, Molkentin JD and Wang X. Proteasome functional insufficiency activates the calcineurin-NFAT pathway in cardiomyocytes and promotes maladaptive remodelling of stressed mouse hearts. Cardiovasc Res. 2010;88:424–33.

35. Aguiar PM, de Mendonca Lima T, Colleoni GWB and Storpirtis S. Efficacy and safety of bortezomib, thalidomide, and lenalidomide in multiple myeloma: An overview of systematic reviews with meta-analyses. Crit Rev Oncol Hematol. 2017;113:195–212.

36. Chang HM, Moudgil R, Scarabelli T, Okwuosa TM and Yeh ETH. Cardiovascular Complications of Cancer Therapy: Best Practices in Diagnosis, Prevention, and Management: Part 1. J Am Coll Cardiol. 2017;70:2536–2551.

37. Fan T, Huang Z, Wang W, Zhang B, Xu Y, Mao Z, Chen L, Hu H and Geng Q. Proteasome inhibition promotes autophagy and protects from endoplasmic reticulum stress in rat alveolar macrophages exposed to hypoxia-reoxygenation injury. J Cell Physiol. 2018;233:6748–6758.

38. Selimovic D, Porzig BB, El-Khattouti A, Badura HE, Ahmad M, Ghanjati F, Santourlidis S, Haikel Y and Hassan M. Bortezomib/proteasome inhibitor triggers both apoptosis and autophagy-dependent pathways in melanoma cells. Cell Signal. 2013;25:308–18.

39. Tannous P, Zhu H, Nemchenko A, Berry JM, Johnstone JL, Shelton JM, Miller FJ, Jr., Rothermel BA and Hill JA. Intracellular protein aggregation is a proximal trigger of cardiomyocyte autophagy. Circulation. 2008;117:3070–8.

40. Kyrychenko VO, Nagibin VS, Tumanovska LV, Pashevin DO, Gurianova VL, Moibenko AA, Dosenko VE and Klionsky DJ. Knockdown of PSMB7 induces autophagy in cardiomyocyte cultures: possible role in endoplasmic reticulum stress. Pathobiology. 2014;81:8–14.

41. Pandey UB, Nie Z, Batlevi Y, McCray BA, Ritson GP, Nedelsky NB, Schwartz SL, DiProspero NA, Knight MA, Schuldiner O, Padmanabhan R, Hild M, Berry DL, Garza D, Hubbert CC, Yao TP, Baehrecke EH and Taylor JP. HDAC6 rescues neurodegeneration and provides an essential link between autophagy and the UPS. Nature. 2007;447:859–63.

42. Schlossarek S, Englmann DR, Sultan KR, Sauer M, Eschenhagen T and Carrier L. Defective proteolytic systems in Mybpc3-targeted mice with cardiac hypertrophy. Basic Res Cardiol. 2012;107:235.

43. Fehling HJ, Swat W, Laplace C, Kuhn R, Rajewsky K, Muller U and von Boehmer H. MHC class I expression in mice lacking the proteasome subunit LMP-7. Science. 1994;265:1234–7.

44. Basler M, Moebius J, Elenich L, Groettrup M and Monaco JJ. An altered T cell repertoire in MECL-1-deficient mice. J Immunol. 2006;176:6665–72.

45. Sha Z, Schnell HM, Ruoff K and Goldberg A. Rapid induction of p62 and GABARAPL1 upon proteasome inhibition promotes survival before autophagy activation. J Cell Biol. 2018;217:1757–1776.

46. Crighton D, Wilkinson S, O’Prey J, Syed N, Smith P, Harrison PR, Gasco M, Garrone O, Crook T and Ryan KM. DRAM, a p53-induced modulator of autophagy, is critical for apoptosis. Cell. 2006;126:121–34.

47. Kenzelmann Broz D, Spano Mello S, Bieging KT, Jiang D, Dusek RL, Brady CA, Sidow A and Attardi LD. Global genomic profiling reveals an extensive p53-regulated autophagy program contributing to key p53 responses. Genes Dev. 2013;27:1016–31.

48. Jain A, Lamark T, Sjottem E, Larsen KB, Awuh JA, Overvatn A, McMahon M, Hayes JD and Johansen T. p62/SQSTM1 is a target gene for transcription factor NRF2 and creates a positive feedback loop by inducing antioxidant response element-driven gene transcription. J Biol Chem. 2010;285:22576–91.

49. Copetti T, Bertoli C, Dalla E, Demarchi F and Schneider C. p65/RelA modulates BECN1 transcription and autophagy. Mol Cell Biol. 2009;29:2594–608.

50. Ling J, Kang Y, Zhao R, Xia Q, Lee DF, Chang Z, Li J, Peng B, Fleming JB, Wang H, Liu J, Lemischka IR, Hung MC and Chiao PJ. KrasG12D-induced IKK2/beta/NF-kappaB activation by IL-1alpha and p62 feedforward loops is required for development of pancreatic ductal adenocarcinoma. Cancer Cell. 2012;21:105–20.

51. Trivedi PC, Bartlett JJ, Perez LJ, Brunt KR, Legare JF, Hassan A, Kienesberger PC and Pulinilkunnil T. Glucolipotoxicity diminishes cardiomyocyte TFEB and inhibits lysosomal autophagy during obesity and diabetes. Biochim Biophys Acta. 2016;1861:1893–1910.

52. Li HH, Kedar V, Zhang C, McDonough H, Arya R, Wang DZ and Patterson C. Atrogin-1/muscle atrophy F-box inhibits calcineurin-dependent cardiac hypertrophy by participating in an SCF ubiquitin ligase complex. J Clin Invest. 2004;114:1058–71.

53. Wang W, Gao Q, Yang M, Zhang X, Yu L, Lawas M, Li X, Bryant-Genevier M, Southall NT, Marugan J, Ferrer M and Xu H. Up-regulation of lysosomal TRPML1 channels is essential for lysosomal adaptation to nutrient starvation. Proc Natl Acad Sci U S A. 2015;112:E1373–81.

54. Zhang X, Cheng X, Yu L, Yang J, Calvo R, Patnaik S, Hu X, Gao Q, Yang M, Lawas M, Delling M, Marugan J, Ferrer M and Xu H. MCOLN1 is a ROS sensor in lysosomes that regulates autophagy. Nat Commun. 2016;7:12109.

55. Perez-Galan P, Roue G, Villamor N, Montserrat E, Campo E and Colomer D. The proteasome inhibitor bortezomib induces apoptosis in mantle-cell lymphoma through generation of ROS and Noxa activation independent of p53 status. Blood. 2006;107:257–64.

56. Turco E, Fracchiolla D and Martens S. Recruitment and Activation of the ULK1/Atg1 Kinase Complex in Selective Autophagy. J Mol Biol. 2019.

57. Cha-Molstad H, Yu JE, Feng Z, Lee SH, Kim JG, Yang P, Han B, Sung KW, Yoo YD, Hwang J, McGuire T, Shim SM, Song HD, Ganipisetti S, Wang N, Jang JM, Lee MJ, Kim SJ, Lee KH, Hong JT, Ciechanover A, Mook-Jung I, Kim KP, Xie XQ, Kwon YT and Kim BY. p62/SQSTM1/Sequestosome-1 is an N-recognin of the N-end rule pathway which modulates autophagosome biogenesis. Nat Commun. 2017;8:102.

58. Sanchez-Martin P and Komatsu M. p62/SQSTM1 - steering the cell through health and disease. J Cell Sci. 2018;131.

59. Duran A, Amanchy R, Linares JF, Joshi J, Abu-Baker S, Porollo A, Hansen M, Moscat J and Diaz-Meco MT. p62 is a key regulator of nutrient sensing in the mTORC1 pathway. Mol Cell. 2011;44:134–46.

