## Supplemental Table and Figures for "The Calcineurin-TFEB-p62 Pathway Mediates the Activation of Cardiac Macroautophagy by Proteasomal Malfunction"

Bo Pan, et al.

### Online Supplements

I. Supplementary Tables 1 and 2
II. Supplementary Figures 1 through 8

### I. Supplementary Tables

| Supplementary Table 1. Antibodies Used in This Study | | | | |
|---|---|---|---|---|
| Antibodies for | Source | Catalog # | Dilution for western blot | Dilution for immunofluorescence |
| α-actinin | Sigma-Aldrich | A5228 | 1:3000 | |
| GAPDH | Sigma-Aldrich | G8795 | 1:1000 | |
| GFP | Santa Cruz | sc-9996 | 1:3000 | |
| Histone H3 | Sigma | H0164 | 1:1000 | |
| LC3 | Cell Signaling | 2775 | 1:2000 | |
| PSMC1 | Enzo Life Sciences | PW0530 | 1:1000 | 1:100 |
| p62 | Cell Signaling | 5114 | 1:1000 | |
| p62 | American Research Products | 03-GP62-C | 1:1000 | 1:500 |
| TFEB | Bethyl Laboratories | A303-673A | 1:1000 | 1:200 |
| Tubulin | Developmental Studies Hybridoma Bank | E7 | 1:5000 | |
| ubiquitin | Cell Signaling | 3936S | 1:1000 | 1:100 |
| Alexa Fluor®-488 | Abcam | ab150073 | | |
| Alexa Fluor® 488 Goat Anti-Rabbit IgG (H+L**)** | Jackson ImmunoResearch | 111-545-003 | | 1:500 |
| Alexa Fluor™ 568 Phalloidin | Thermo Fisher Scientific | A12380 | | 1:100 |

| Supplementary Table 2. PCR Primers | | | |
|---|---|---|---|
| **Rat genes** | **Primers (5' – 3')** | **Mouse genes** | **Primers (5' – 3')** |
| Becn1 | TTACTTACCACAGCCCAGG; TGCTCCAACATCTCCAAAC | M6PR | CTCATCTCCCTCTACTGGTACTT; CGCTTCCATTCCCTTCCTTTA |
| CtsB | TGGGTTCAGCGAGGACAT; ATGGTGTAGGGTAAGCAGCC | Mcoln1 | CTGTCATCTACCTGGGCTATTG; GAGTGAGAACAGACACTCAGAAA |
| CtsD | GCGTCTTGCTGCTCATTCT; AACTCTGACACTGGCTCCTT | p62 | CTCTGGACACGATCCAGTATTC; CTGCTCTACGTGATGCAACTA |
| GAPDH | ATGACATCAAGAAGGTGGTG; CATACCAGGAAATGAGCTTG | Psmc1 | TCAGCATCCTGTCGTTTGTAG; GGGATCCGTGTCATCCATTAG |
| Lamp1 | GTGGGACTTGCGGTGCC; GACATTGAGGGCGAGCG | Uvrag | GACCACGAGACAGTTGAGATAG; GCAGGGACAATGGACTTAGAA |
| M6PR | GGATAAGGAGTCAAAGAATG; TGATTCTCCCAACCACCGT | Vps18 | GGACTTGATGGCTTTGTGTTG; CGTGACCTGACCTGTTCTATTC |
| MCIP1.4 | AGCTCCCTGATTGCCTGTGT; TTTGGCCCTGGTCTCACTTT | | |
| Mcoln1 | CGCCGCCGCCTCAAGT; GCTGCTCCCGTGTGTAGGC | | |
| p62 | GCCACCTCTCTGATAGC; AGGTTTGCTGACTTCCG | | |
| Rab7a | CAGTCTCTTGGTGTGGC; CGAAGTAAGGAATGTTG | | |
| Uvrag | CTTCTGGATACCTACTTCAC; GACTTTCCACTCTATCAACAGC | | |
| Vps18 | GCTCCGCATTGACTTGGG; GCCTTCTGTCCATTGCGGT | | |

### II. Supplementary Figures

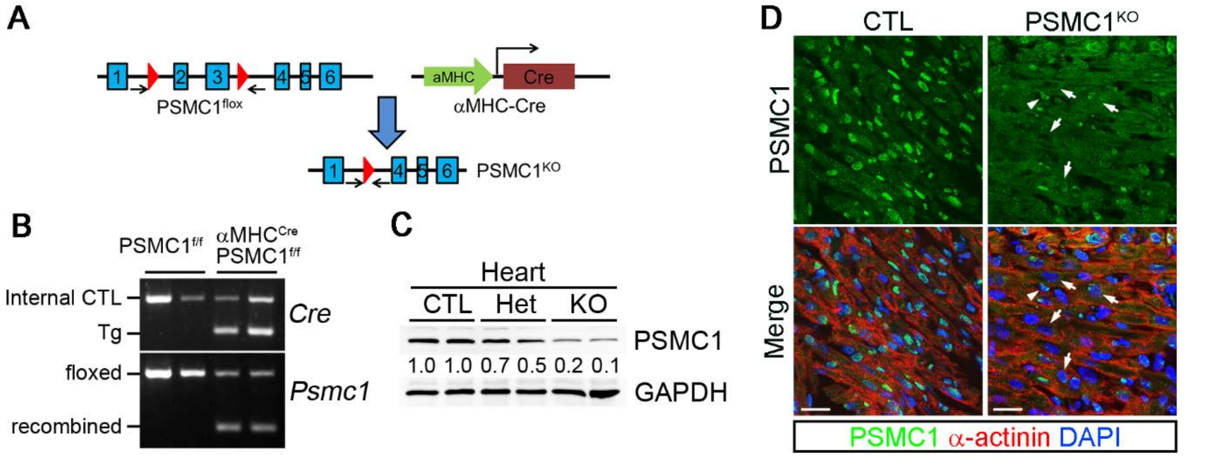

**Supplementary Figure 1. Generation of cardiomyocyte-restricted Psmc1 knockout (Psmc1$^{CKO}$) mice. A**, Schematic diagram depicting the strategy of **Psmc1$^{CKO}$** in cardiomyocytes. Arrows denote the binding sites of the primers that are used to detect the presence and recombination of floxed *Psmc1* alleles. **B**, PCR genotyping. **C**, Western blots of indicated proteins in the hearts of 2-day-old CTL (PSCM1$^{f/f}$), heterozygous (Het, αMyHC-Cre:PSCM1$^{f/+}$) and KO (αMyHC-Cre:PSCM1$^{f/f}$) mice. The relative abundance of Psmc1 proteins was marked below the blot. **D**, Confocal images of immunostained myocardium sections from P2 mice. Psmc1 (green) is nuclear-enriched in CTL hearts but this staining pattern was lost in the cardiomyocytes of KO hearts (arrows). The arrowhead points to a non-cardiomyocyte that retains the nuclear-enriched Psmc1 staining in KO hearts. The sections were counterstained with α-actinin and DAPI, respectively. Bar, 20 µm.

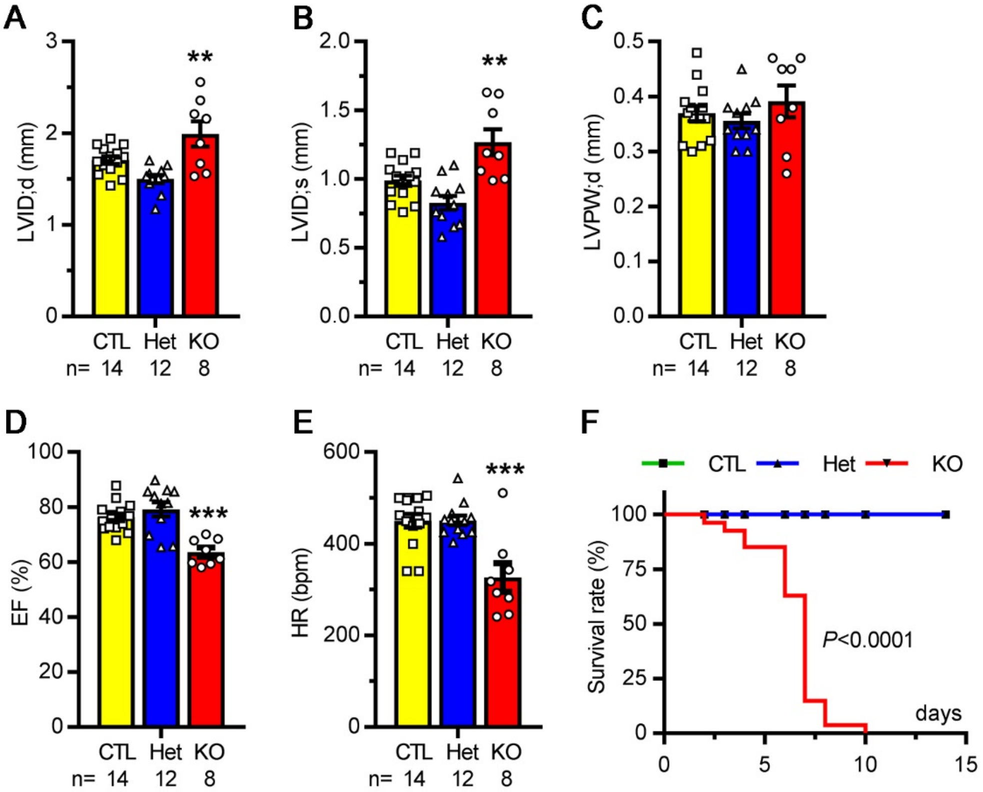

**Supplementary Figure 2**. **Psmc1$^{CKO}$ caused heart failure and perinatal lethality**. **A-E,** Echocardiographic analysis of Psmc1$^{CKO}$ mice. Two-day-old mice were gently secured on the station with tapes and kept conscious during the procedure. Cardiac images were recorded using a VEVO 770 echocardiography system with a 30MHz transducer (Visual Sonics). The LV morphometric and functional parameters were analyzed off-line using VEVO 770 software. The LV internal diameter at diastole (A) and systole (B), LV posterior wall thickness at diastole (C), LV ejection fraction (EF) and heart rate (HR) are shown. ** P < 0.01, *** P<0.001 vs CTL or Het. **F,** Kaplan-Meier survival analysis. N for CTL, Het and KO are 59, 47 and 27 respectively. All of the KO mice died before P10.

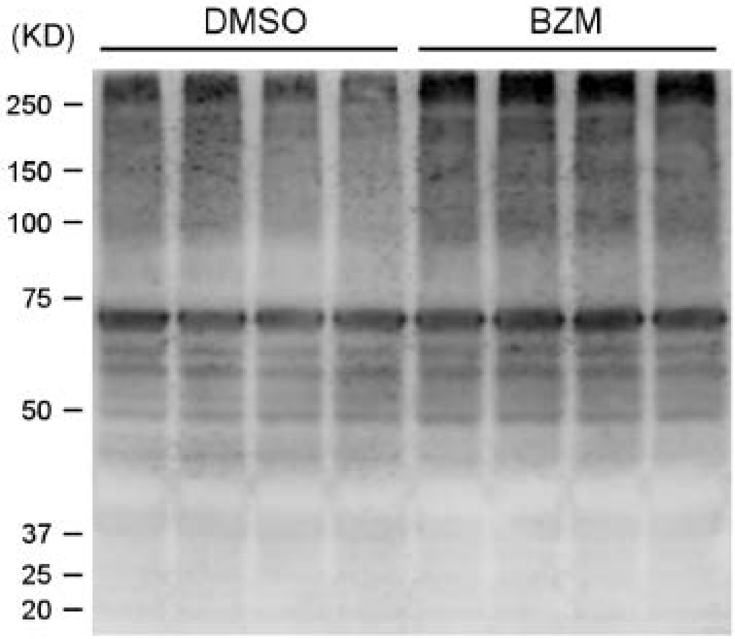

**Supplementary Figure 3**. Western blot analysis for myocardial levels of total ubiquitinated proteins in bortezomib (BZM) treated mice. Mice were treated with BZM (10 ug/kg, i.p.) or vehicle control (60% DMSO in saline) and ventricular myocardium was sampled as described for Figure 3 of main text.

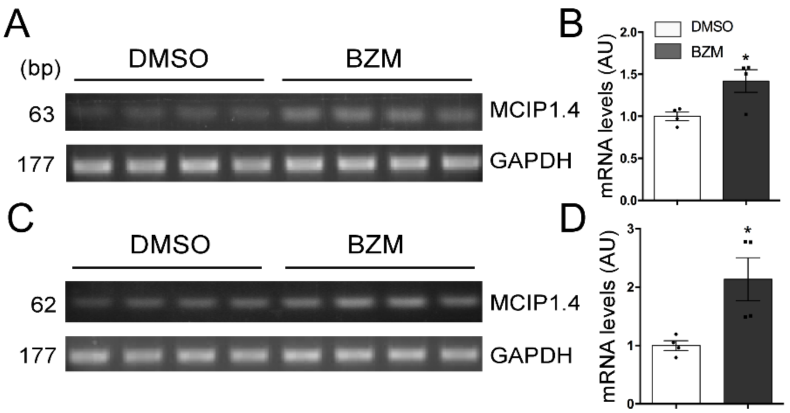

**Supplementary figure 4**. BZM increases MCIP1.4 mRNA levelsin mouse hearts (A, B) and cultured NRVMs (C, D).

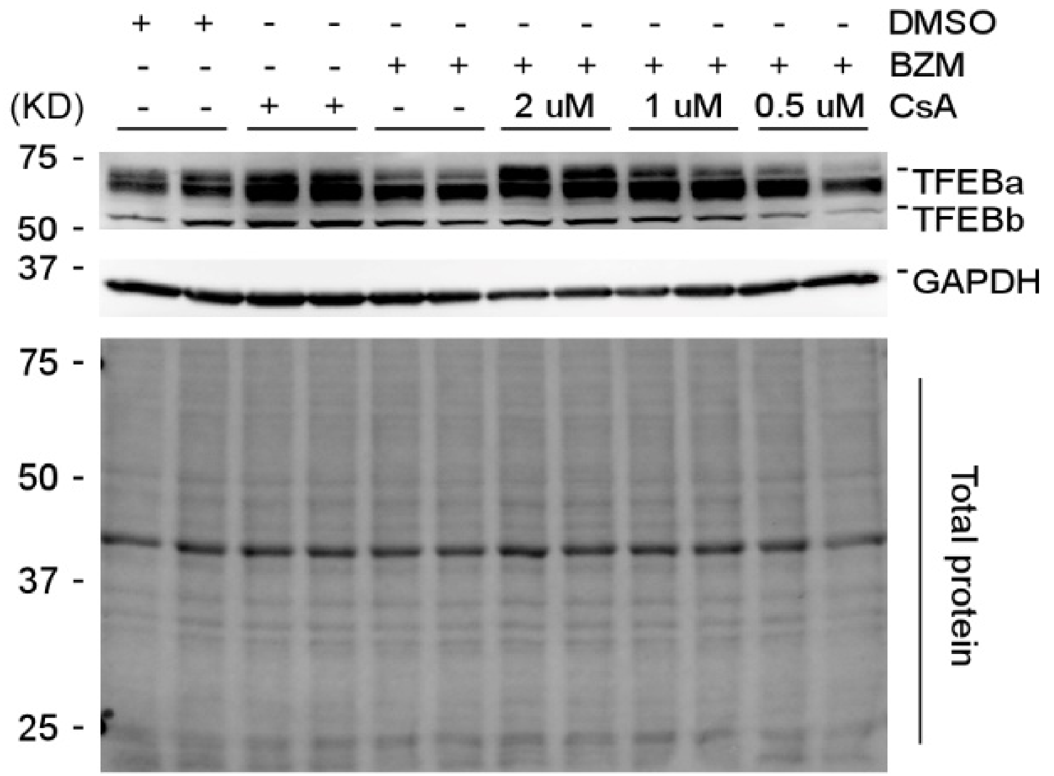

**Supplementary Figure 5.** BZM induction of TFEB dephosphorylation in cultured NRVMs is attenuated by calcineurin inhibition with cyclosporine A (CsA).

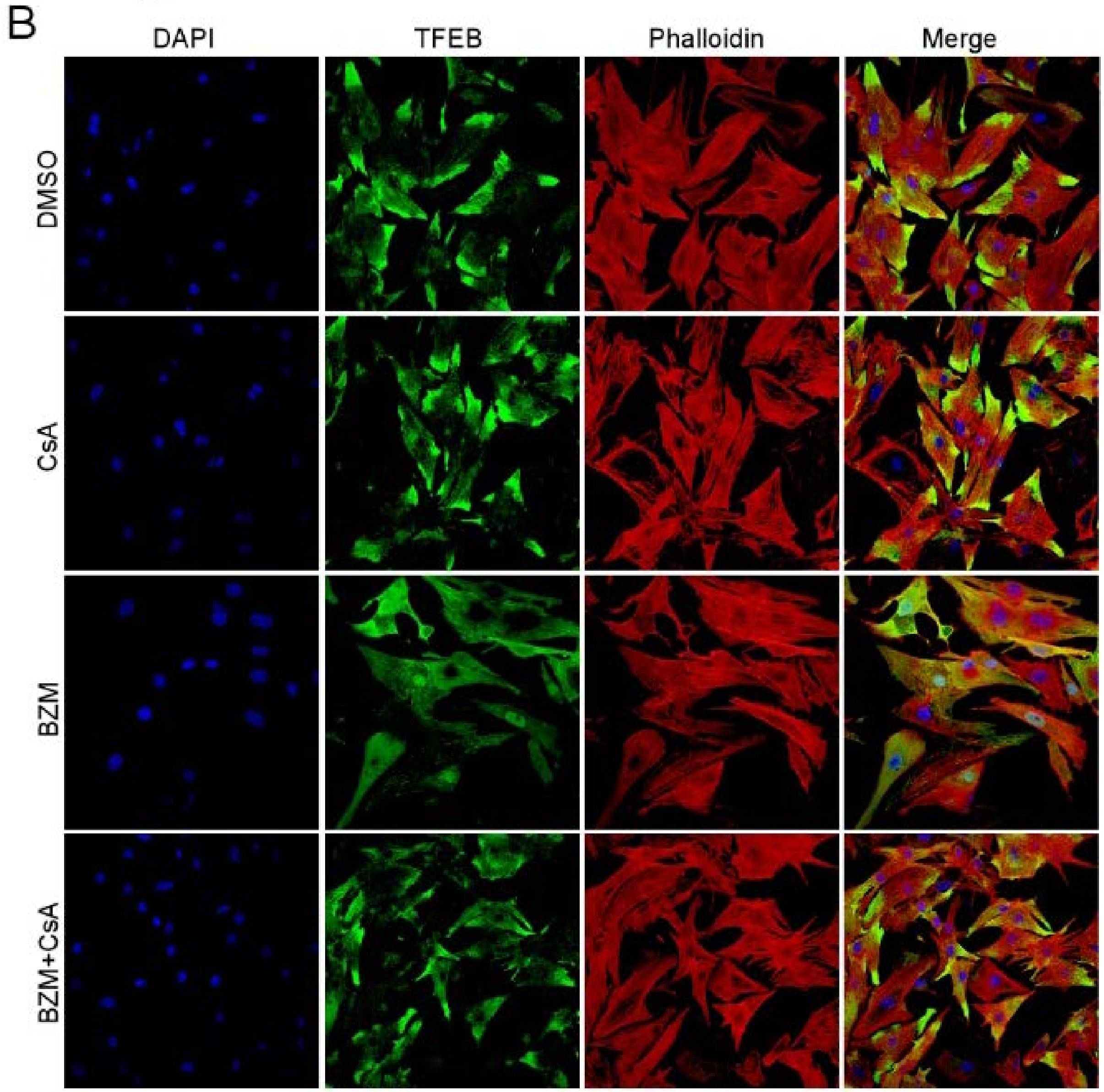

**Supplementary Figure 6.** Increased nuclear localization of TFEB by PSMI in NRVMs is calcineurin-dependent.

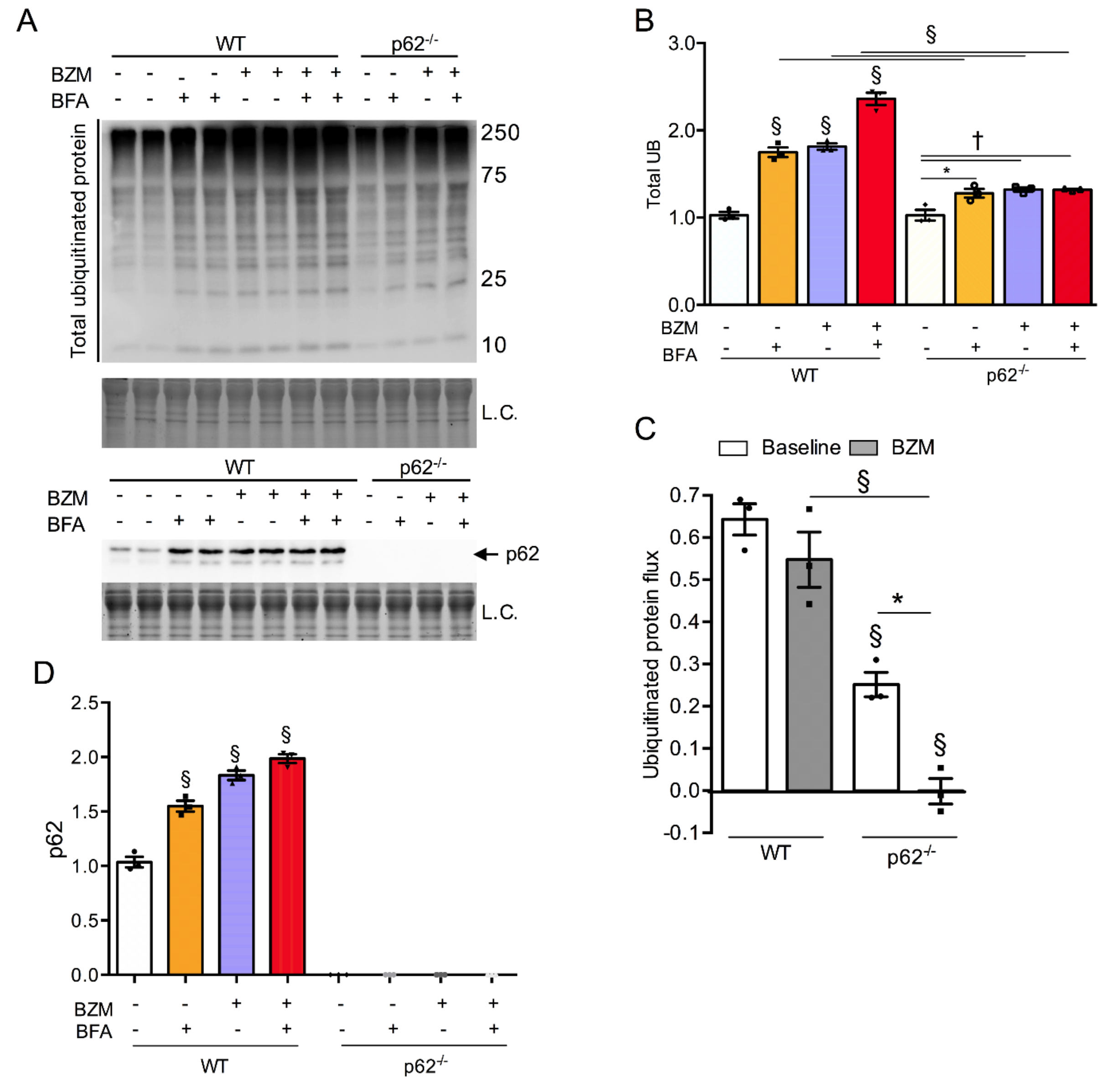

**Supplementary figure 7. p62 deficiency (p62$^{-/-}$) significantly reduces lysosomal degradation of ubiquitinated proteins in cultured neonatal mouse cardiomyocytes subjected to PSMI**. Cardiomyocytes isolated from WT and p62$^{-/-}$ mice at postnatal day 2 were cultured for 48 h before subjected to the indicated treatment. The treatment of proteasome inhibitor bortezomib (BZM; 25 nM) lasted 6 hours before initiating of the treatment with bafilomycin A1 (BFA; 25 nM). The cells were harvested after 6 hours of BFA treatment. Total cell lysates were prepared for western blot analyses of total ubiquitinated proteins and p62. Stain-free total protein imaging technology was used as loading control (L.C.; only a segment of the image is shown) to normalize loading variation. Data were analyzed by two-way (C) or three-way (B, D) ANOVA followed by Tukey's post hoc multiple comparisons tests and are presented as Mean ± SEM; n = 3 mice/group. Symbols above the bars indicate the p value for comparison with the vehicle control; **p*<0.05, †*p*<0.01, ‡*p*<0.001, §*p*<0.0001.

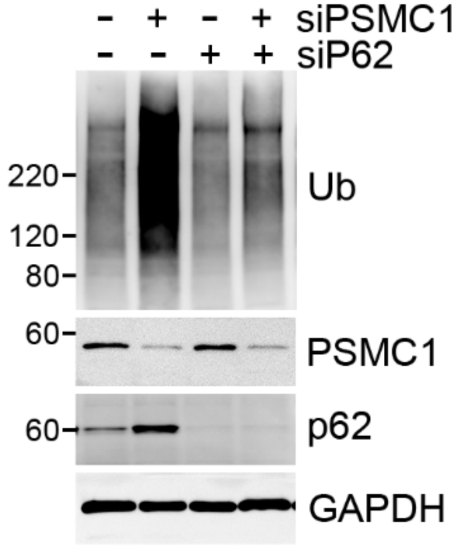

**Supplementary Figure 8. Depletion of p62 diminished the accumulation of ubiquitinated proteins induced by genetic inhibition of proteasome.** NRVMs were transfected with the indicated siRNA for 72 hours. Cell lysates were used for western blot of the indicated proteins. Ub, ubiquitinated proteins.
